# DiVerG: Scalable Distance Index for Validation of Paired-End Alignments in Sequence Graphs

**DOI:** 10.1101/2025.02.12.637964

**Authors:** Ali Ghaffaari, Alexander Schönhuth, Tobias Marschall

## Abstract

Determining the distance between two loci within a genomic region is a recurrent operation in various tasks in computational genomics. A notable example of this task arises in paired-end read mapping as a form of validation of distances between multiple alignments. While straightforward for a single genome, graph-based reference structures render the operation considerably more involved. Given the sheer number of such queries in a typical read mapping experiment, an efficient algorithm for answering distance queries is crucial. In this paper, we introduce DiVerG, a compact data structure as well as a fast and scalable algorithm, for constructing distance indexes for general sequence graphs on multi-core CPU and many-core GPU architectures. DiVerG is based on PairG [28], but overcomes the limitations of PairG by exploiting the extensive potential for improvements in terms of scalability and space efficiency. As a consequence, DiVerG can process substantially larger datasets, such as whole human genomes, which are unmanageable by PairG. DiVerG offers faster index construction time and consistently faster query time with gains proportional to the size of the underlying compact data structure. We demonstrate that our method performs favorably on multiple real datasets at various scales. DiVerG achieves superior performance over PairG; e.g. resulting to 2.5–4x speed-up in query time, 44–340x smaller index size, and 3–50x faster construction time for the genome graph of the MHC region, as a particularly variable region of the human genome.

The implementation is available at: https://github.com/cartoonist/diverg

## 1 Introduction

Many genomic studies use read mapping as a central step in order to place the donor sequence reads into context relative to a reference. Several studies have repeatedly shown that the conventional reference assemblies, i.e. consensus genomes or genomes of individuals, do not capture the genomic diversity of the population and introduce biases [40, 23, 6, 2, 12]. Furthermore, the absence of alternative alleles in a linear reference can penalize correct alignments and lead to decreased accuracy in downstream analysis. To address this issue, augmented references, often in the form of *pangenome graphs*, have been developed by incorporating genomic variants observed in a population [16].

On the other hand, shifting from a linear structure to a graph introduces a variety of theoretical challenges. Recent progress in *computational pangenomics* has paved the way for using such references in practice as established algorithms and data structures for linear references cannot be seamlessly applied to their graphical counterparts [9, 16, 39]. One significant measure affected by this transition is the concept of *distance* between two genomic loci. Defining and computing distance relative to a single genome is inherently straightforward. However, in genome graphs, distance cannot be uniquely defined due to the existence of multiple paths between two loci, with the number of paths being theoretically exponential relative to the number of variants between them.

Determining genomic distances between two loci emerges in several genomic workflows notably paired-end short-read mappings to sequence graphs. In paired-end sequencing, DNA fragments are sequenced from both their ends, where the unsequenced part in between introduces a gap between the sequenced ends. Read mapping is the process of determining the origin of each read relative to a reference genome through sequence alignment. There are several sources of ambiguity in finding the correct alignments. The distance between two pairs plays a crucial role in resolving alignment ambiguities, for example in repetitive regions [5]. An accurate alignment of one end can rescue the other end’s alignment in case of ambiguities. Therefore, it is important to determine whether the distance between two reference loci, where two ends of a read could be placed, falls in a particular, statistically well motivated range [*d*_1_, *d*_2_]. This problem, referred to as the *Distance Validation Problem (DVP)*, is first formally defined in [28].

The distance between all pairs of candidate alignments corresponding to a paired-end reads should be validated in order to find the correct pairs. Additionally, the candidate alignments in pangenome graphs are often more abundant compared to a linear sequence due to the increased ambiguity in a pangnome reference caused by added variations. This fact, combined with the large number of reads in a typical read mapping workflow, necessitates efficient methods to answer the DVP, often by preprocessing the graph and constructing an index data structure. Therefore, the efficiency of the involved operation is crucial for rendering short-read-to-graph mapping practically feasible.

Long, third-generation sequencing reads (TGS) have spurred enormous enthusiasm in various domains of application, which may explain that the majority of read-to-graph mapping approaches focuses on such long reads. However, still, long TGS reads are considerably more expensive to produce than short reads.

Short next-generation sequencing reads are highly accurate, cheap, and available to nearly every sequencing laboratory today. This provides substantial motivation for delivering approaches that render short-read-to-graph mapping a viable option; in fact, this would free the way for using graph-based reference systems in many laboratories worldwide and enable their use for existing biobank-scale cohorts where short reads have already been produced.

### Related Work

The distance between two nodes in a graph is determined by the paths that connect them and is typically associated with their shortest path. Identifying the shortest paths between two points in a graph is a prevalent problem across various application domains and is an extensively studied topic in computer science [14, 18, 19, 25, 27, 41, 15]. However, simply knowing the shortest path length is not always sufficient for addressing the DVP: when the shortest path length is less than *d*_1_, it cannot determine whether a longer path of length *d* exists satisfying *d*_1_≤ *d* ≤*d*_2_. On the other hand, the decision version of *longest path problem* and *exact-path length problem*, which respectively answer whether there is a path of at least length *d* or exactly length *d* in a graph, are shown to be NP-complete [38, 33].

In the context of sequence graphs, most sequence-to-graph aligners use heuristic approaches to estimate distances [21, 36]. Chang et al. [7] propose an exact method and indexing scheme to determine the minimum distance between any two positions in a sequence graph with a focus on seed clustering. Although their method is shown to be efficient in seed clustering, it cannot address the DVP as it only provides the minimum distance between two positions.

Jain et al. [28] propose PairG, a distance indexing method which, to the best of our knowledge, is the first method directly addressing the DVP. This method, similar to existing approaches, involves preprocessing the graph to create an index that answers distance queries efficiently. In PairG, the index data structure is a sparse Boolean matrix constructed from powers of the adjacency matrix of the graph. Once constructed, it can indicate in near-constant time whether a path exists between two nodes that meet the distance criteria. It benefits from general sparse storage format and employs standard sparse matrix algorithms to reduce space requirement and accelerate the index construction. However, the method cannot scale to handle whole genome graphs for large sequences. PairG, although applicable for small graphs, also does not run on many-core architectures such as GPU for sparse matrix computations due to its intense memory requirements.

### Contributions

In this work, we propose DiVerG, an indexing scheme that offers fast exact solutions for the DVP in sequence graphs with significantly lower memory footprint and faster query and construction time than existing methods.

DiVerG enhances PairG to overcome the limitations of existing approaches. Our first contribution is new *dynamic* compressed formats, namely *rCRS* formats, for storing sparse Boolean matrices. They are *dynamic* in the sense that sparse matrix operations can be conducted directly on matrices in this format without decompression. These formats, although simple by design, provide considerably greater compression potential for sparse matrices representing the adjacency matrix of sequence graphs, or the powers thereof. The specific incorporation of the logic that supports the shape of adjacency matrices of sequence graphs facilitates significantly more compact representations than what can be achieved by merely utilizing the sparsity of the matrix in standard sparse formats.

Secondly, we propose algorithms for Boolean sparse matrix-matrix multiplication and matrix addition for matrices in rCRS formats. Our multiplication algorithm achieves enhanced performance through the bit-level parallelism that the encoded format immediately provides, while the addition algorithm runs in time proportional to the compressed size.

We tailored both algorithms and their implementations to be particularly powerful on massively parallel architectures, such as GPUs. The faster algorithms and more compacted encodings enable indexing of drastically larger graphs, previously infeasible with the prior state of the art. Our experiments show that DiVerG scales favorably with large sequence graphs at low memory footprint and significant compression ratio. Most importantly, DiVerG responds to distance queries in constant time in practice. Despite the prior work, DiVerG’s query performance does not grow by the distance constraints parameters.

## 2 Problem Definition

### 2.1 Sequence Graphs

Sequence graphs provide compressed representations of sets of similar – evolutionary or environmentally related – genomic sequences [26]. They are typically defined as node-labelled *bidirected* graphs where each node contains a string label. Bidirectionality allows traversal in both forward and reverse directions, which accounts for the complementary structure of genomes. For theoretical analysis, we use an equivalent representation called a *character graph*, defined as *G*(*V, E, λ*) where *V* and *E* are the node and edge sets, respectively, and *λ* : *V*→ Σ assigns a *single* character from alphabet Σ to each node. Character graphs and general sequence graphs can be converted to each other in time linear in the size of the character graph [28]. By definition, in a character graph, the length of a sequence spelt out by a walk of length *k* is exactly *k*. This one-to-one correspondence between walk length and sequence length is key to the theoretical foundation of our method.

Lastly, a *chain graph* is a character graph that represents a single sequence which is equivalent to the sequential chaining of all (single-letter) nodes. A sequence graph *G* is called *sparse* if the average node degree in *G* is close to 1.

### 2.2 Distance Validation Problem

Fragments in sequencing libraries typically vary in size depending both on the sequencing technology in use and library preparation procedures. In our study, the distance between paired reads in a library is modelled by an interval indicating the expected lower (*d*_1_) and upper bounds (*d*_2_). This interval, which is referred to as *distance constraints* and denoted as (*d*_1_, *d*_2_), are assumed to be provided as input parameters. In Appendix A, we discussed how the distance constraints can be determined for a library.

As stated before, aligning short reads to a reference genome, whether linear or graph, can be ambiguous due to several factors such as repetitive regions in the genome, sequencing errors, and the genetic differences between the donor genome and the reference. In such cases, reads might have multiple candidate alignments. Utilizing the distance information in paired-end sequencing data can help identify the correct alignments (Figure 9b in Appendix A).

Two alignments are considered as paired if the the distance between them is consistent with the inferred fragment model. In addition to the distance, the orientation of two paired reads should also be as expected depending on the sequencing technology in use; e.g. one of the pair should be aligned on the forward strand while the other is on the reverse strand.

#### Problem 1

(Distance Validation). *Let r*_1_ *and r*_2_ *be two paired reads, and a*_1_, *and a*_2_ *their alignments against a sequence graph G*(*V, E, λ*). *Having a*_1_ *mapped to the forward strand implies a*_2_ *to be on the reverse strand. Let v*_1_ *and v*_2_ *be the first nodes in the paths to which a*_1_ *and a*_2_ *are aligned, respectively (Figure 1). Assuming the fragment model is described by distance constraints* (*d*_1_, *d*_2_), *the Distance Validation problem is determining whether there exists a path from v*_1_ *to v*_2_ *of length d* ∈ [*d*_1_, *d*_2_]. *These two alignments are considered as* paired *if this condition is met*.

**Figure 1.**
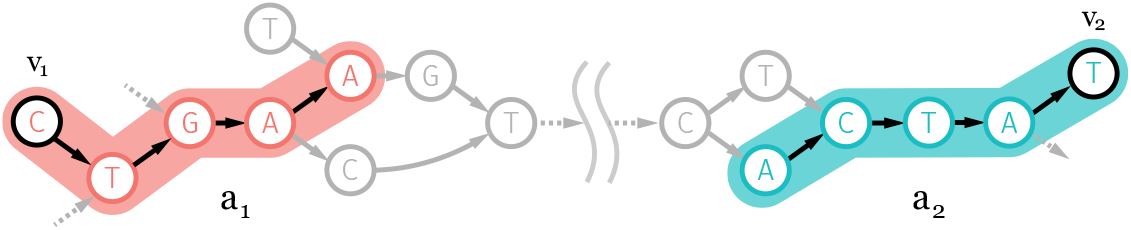
Alignments *a*_1_ and *a*_2_ correspond to paired reads *r*_1_ = CTGAA and *r*_2_ = ATAGT on a sequence graph (partially represented). Alignment *a*_1_ starts from node *v*_1_ and extends along the forward strand, while alignment *a*_2_ starts from *v*_2_ and extends along the reverse complement strand.

Due to the sheer number of queries in a typical read mapping experiment, the algorithm solving Problem 1 needs to be efficient.

## 3 *k*-Walk Matrix as Distance Index

Let **A** = (*a*_*ij*_) denote the Boolean adjacency matrix of graph *G*(*V, E, λ*), defined by *a*_*ij*_ = 1 if and only if (*i, j*) *∈ E*. The *k−*th power of **A** has a special property: 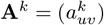 determines the number of walks of length *k* between nodes *u* and *v* in *G*. The Boolean equivalent of **A**^*k*^, which can be achieved by replacing each non-zero entry in **A**^*k*^ with 1, can answer *existence* queries on walks of length *k* instead.

For the remainder of the paper, any mentioned adjacency matrices or their *k*-th powers implicitly refer to Boolean matrices.

### Definition 2

(Boolean Matrix Operations). *Given two Boolean square matrices* **A** *and* **B** *of order n, standard Boolean matrix operations are defined as follows:*

▂ *Addition:* **A** ∨ **B** = *a*_*ij*_ ∨ *b*_*ij*_,
▂ *Multiplication:* 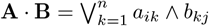
▂ *Power:* **A**^*k*^ = **A** *·* **A**^*k−*1^ (*k >* 0), *and* **A**^0^ = **I**,

*where* ∨ *and* ∧ *are Boolean disjunction (or) and conjunction (and) operators*.

### Definition 3

(Distance Index). *Given distance constraints* (*d*_1_, *d*_2_), *a Boolean matrix 𝒯 is called* distance index *relative to* (*d*_1_, *d*_2_) *if defined as:*

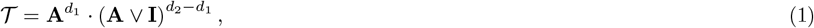

where *I* is the identity matrix, and denote Boolean matrix addition and matrix-matrix multiplication, respectively, defined in Definition 2. Jain et al. [28] show that 𝒯 = (*τ*_*ij*_) can efficiently solve the *Distance Validation Problem* defined in Problem 1, i.e. if *τ*_*uv*_ = 1, there exists at least one walk of length *d* ∈ [*d*_1_, *d*_2_] from node *u* to *v* in the graph.

Generally, the diameter of sequence graphs is very large in practice, and the number of edges is on the order of the number of nodes, i.e. | *E*| ~|*V*|. This implies that most sequence graphs exhibit sparse adjacency matrices. Therefore, given that distance constraints are considerably smaller than the graph, i.e. (*d*_2_ *− d*_1_) ≪ |*V*|, one can expect 𝒯 to be sparse. The common approaches for computing Equation (1) leverage the sparsity of the matrices, aiming to operate at time and space complexities proportional to the number of non-zero elements. Sparse matrices are typically stored in sparse formats such as the *Compressed Row Storage* (CRS) [4] and the calculation of Equation (1) relies on the sparse matrix-matrix multiplication (SpGEMM) and sparse matrix addition (SpAdd) algorithms [4].

### Definition 4

(Compressed Row Storage). *Given Boolean squared matrix* **A** *with n rows and nnz*(**A**) *number of non-zero elements, the* Compressed Row Storage *or* Compressed Sparse

Row *format of* **A** *is a row-based representation consisting of two one-dimensional arrays* (*C, R*); *where*

▂ *C* (column indices), *of size nnz*(**A**), *stores the column indices of non-zero elements in* **A** *in row-wise order;*
▂ *R* (row map), *of length n* + 1, *is defined such that, for each row index I* ∈ [0, *n−* 1], *R*(*i*) *gives the index in C where the column indices for row i begin. The final entry in R is set to* |*C*|, *i*.*e. R*(*n*) = *nnz*(**A**).

### Remark 5.

Row map array *R* defines the boundaries of each row in the column index array *C*: the column indices of non-zero entries of row *i* are located in the half-open interval [*R*(*i*), *R*(*i* + 1)) of *C*. This partitions *C* into consecutive subsequences *C* = *C*_0_*C*_1_ *· · · C*_*n−*1_, referred to as the *row decomposition* of *C*, where *C*_*i*_ = [*C*(*R*(*i*)) *· · · C*(*R*(*i* + 1) *−* 1)].

By definition, array *C* does not need to be sorted within each block *C*_*i*_ in CRS format. When the entries of each row in *C* are sorted, we refer to this as *sorted CRS*. Figure 2a demonstrate a toy example of a Boolean matrix in (sorted) CRS format. The required space for storing sparse matrix **A**_*n×m*_ in CRS format is Θ(*n* + nnz(**A**)).

**Figure 2.**
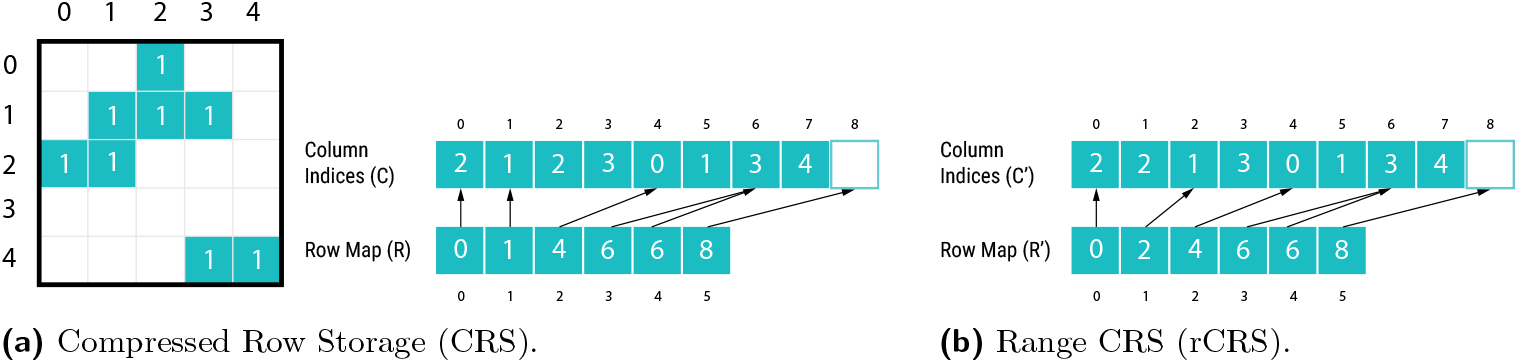
An example Boolean matrix represented in CRS and rCRS format.

Accessing element *a*_*ij*_ in **A** using CRS representation can be reduced to searching for column index *j* in the part of array *C* corresponding to row *i*, as specified by row map array *R*. That is, one searches for *j* in *C*_*i*_, the *i*-th block of the row decomposition of *C*. This search can be facilitated in sorted CRS via binary search, instead of inspecting all column indices in the row. With *z* being the maximum number of non-zero values in any row in **A**, searching for *j* in sorted CRS takes *O*(log(*z*)), due to the binary search performed on the sorted entries. As mentioned before, given the sparsity of matrix **A**, which implies that *z* is very small, accessing elements in sorted-CRS amounts to requiring constant time in practice. To date, many algorithms have been proposed for parallel sparse matrix-matrix multiplication in formats such as CRS or other similar variants [35, 3]. Since standard matrix operations establish fundamental components of numerous applications, most of them are optimized for various hardware architectures, in particular for architectures that support massive parallelization, such as GPUs. Although the CRS format is often sufficient for general sparse matrix storage and relevant operations, it is computationally prohibitive when it is used for computing the *k*-th power of adjacency matrices as well as the resulting matrix from Equation (1) for large sequence graphs. On the other hand, sequence graph adjacency matrices offer optimization opportunities that are not typically found in general sparse matrices.

### 3.1 Observations

The distance index 𝒯 computed by Equation (1) quickly becomes infeasible for large graphs due to its space requirements. For example, consider a human pangenome graph constructed using the autosomes of the GRCh37 reference genome and incorporating variants from the 1000 Genome Project. Constructing the distance index 𝒯 for this graph with distance criteria *d*_1_ = 150 and *d*_2_ = 450 results in a matrix of order 2.8B with approximately 870B non-zero values. Assuming that each non-zero value consumes 4 bytes, the total space required for the final distance matrix in sorted CRS would be about 3.5 TB.

Even for smaller graphs, storage requirements are impractical for GPU architectures as they have much smaller memory than the accessible main memory in a CPU. Such many-core architectures are beneficial for faster computation of SpGEMM, which is the computational bottleneck for constructing the distance index.

A key observation for identifying the compression potential is the structure of the distance matrix. Genome graphs usually consist of multiple connected components corresponding to each genomic region or chromosome. If the nodes are indexed such that their indices are *localized* for each region, the distance index 𝒯 is a *block-diagonal matrix*, where each block corresponds to a different genomic region or chromosome.

Further examinations reveal that in each row, non-zero values are also localized within a certain range – often grouped into a few clusters of consecutive columns – if nodes are indexed in a “near-topological order”. This is because almost all edges in sequence graphs are local, and it is reflected in the matrix as long as the ordering that defines node indices mostly preserves this locality – i.e. adjacent nodes appear together in the ordering. This observation can be seen clearly in the distance index constructed for a chain graph with distance constraints (*d*_1_, *d*_2_) as shown in Figure 3.

**Figure 3.**
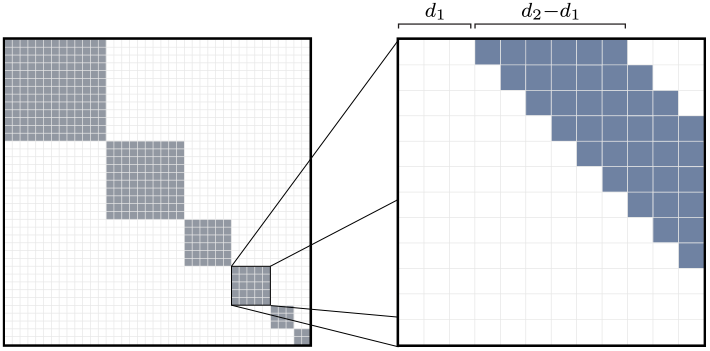
Schematic illustration of the structure of matrix 𝒯 with distance constraints (*d*_1_, *d*_2_) for a chain graph with multiple components.

## 4 Method

### 4.1 Total Order on Node Set

The nodes of a graph are indexed along the rows of its adjacency matrix with the same order governing the columns as well. This induces a total order on the set of nodes in the graph, assigning each node a unique *index*. Different total orders yield different adjacency matrices. As will be shown in Section 4.2, our method greatly benefits if the total order on the node set maximizes the *locality* of the indices for adjacent nodes; i.e. adjacent nodes are assigned nearby indices by such ordering. It can be shown that this problem is equivalent to finding a sparse matrix with minimum bandwidth by permuting its rows and columns. The problem of finding such ordering has been proven to be NP-Complete [34].

Several heuristic algorithms have been proposed to minimize the bandwidth of sparse matrices by reordering rows and columns [10, 11, 8]. However, applying these methods to our problem presents some key limitations. First, although our method logically views sequence graphs as character graphs, in practice it works with string-labelled graphs for space efficiency. This requires preserving the sequential ordering of bases within nodes of a string-labelled graph, a constraint that existing heuristics do not necessarily meet when applied to the character graph representation. Breaking this intra-node ordering introduces memory overhead and fragmentation, impacting graph traversal performance in downstream tasks through reduced memory locality and increased cache misses. Second, when applied directly to string-labelled graphs while preserving node ordering, these methods are less effective at reducing bandwidth compared to their performance on character graphs.

To address these limitations, we propose a new heuristic algorithm inspired by the Cuthill–McKee (CM) algorithm [10, 11]. The new algorithm directly operates on string-labelled sequence graphs. In essence, the algorithm traverses the graph in a breadth-first search similar to the CM. However, instead of prioritizing visiting nodes by their degrees, it prioritizes nodes that have a predecessor with a lower *topological sort order* (Algorithm 1). Note that, in practice, sequence graphs, particularly variation graphs, are DAGs in the majority of relevant cases. If input sequence graphs are not acyclic, e.g. in de Bruijn graphs, we resort to a “semi-topological ordering” established by “dagifying” the graph, i.e. by running topological sort algorithm on the graph while the traversal algorithm ignores any edges forming a cycle.

Our experimental results (in Section 5) demonstrate that the ordering achieved by Algorithm 1 offers a very effective heuristic while assigning sequential indices to bases within each node. This can be justified by the fact that sequence graphs generally preserve the linear structure of the sequences from which they are constructed, and the variations in these sequences are often only local in the genomic coordinates, thereby affecting the graph topology locally. As a result, prioritizing sibling nodes for index assignment while following topological ordering tends to localize indices of adjacent nodes, thereby minimizing bandwidth.

#### Algorithm 1

A heuristic for defining a total order on the node set that minimizes bandwidth.

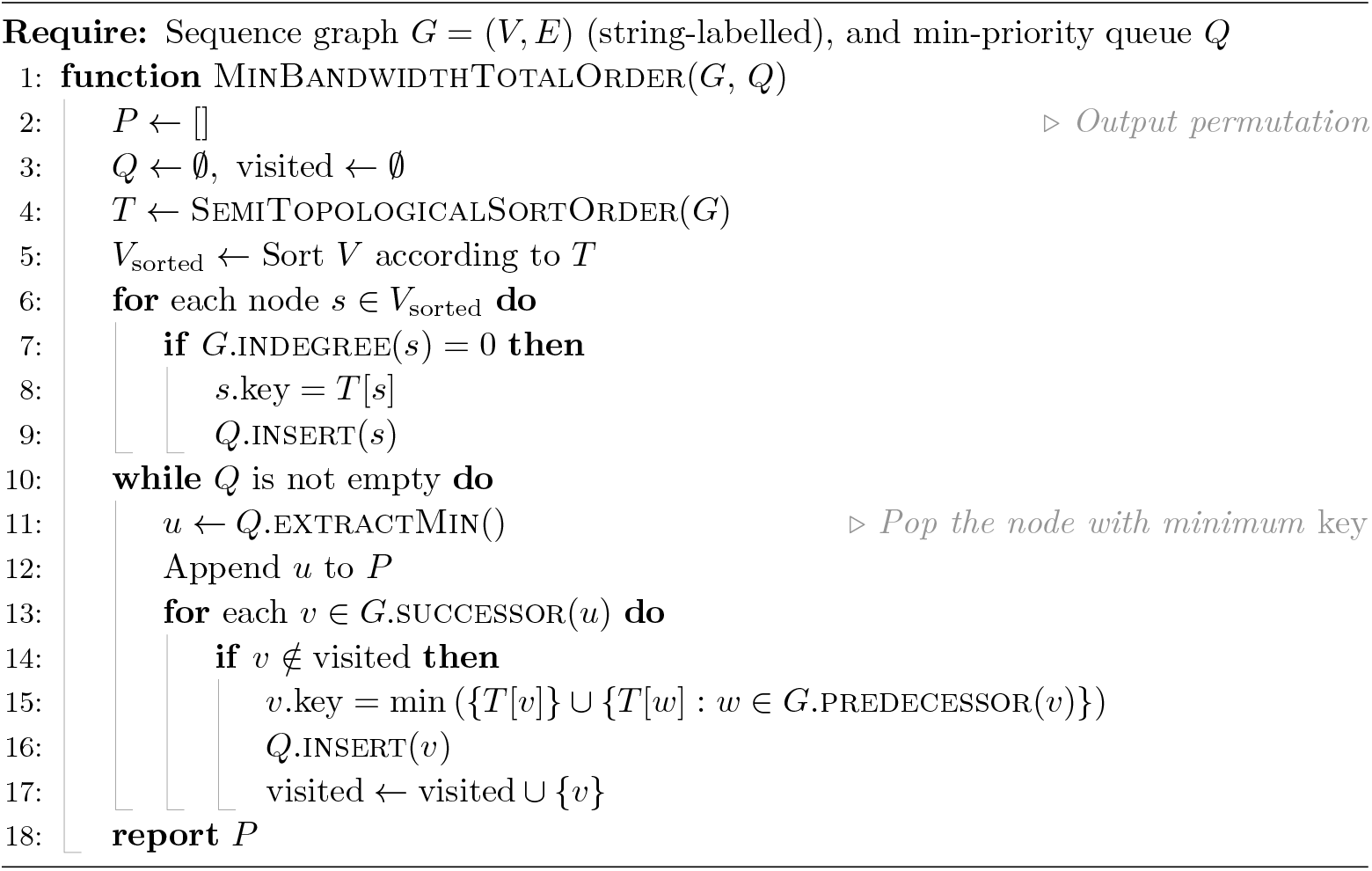

### 4.2 Range Compressed Row Storage (rCRS)

DiVerG aims to exploit particular properties of the distance index matrix 𝒯 mentioned in Section 3.1, as well as its sparsity, to reduce space and time requirements. Our method introduces a new sparse storage format, which will be explained subsequently. This format relies on transforming the standard CRS by replacing column index ranges with their lower and upper bounds to achieve high compression rate. Later, we will propose tailored algorithms to compute sparse matrix operations directly on this compressed data structure without incurring additional overhead for decompression.

#### Definition 6

(Minimum Range Sequence). *Let A be a* sorted *sequence of distinct integers. A* “range sequence” *of A is another sequence, A*_*r*_, *constructed from A in which any disjoint subsequence of consecutive integers is replaced by its first and last values. When such subsequence contains only one integer i, it will be represented as* [*i, i*] *in A*_*r*_. *The minimum sized A*_*r*_ *is defined as* minimum range sequence *of A, denoted as ρ*(*A*).

#### Example 7.

Consider *A* = [10, 11, 12, 13, 23, 29, 30]. It has three disjoint sequence of consecutive integers: 10–13, 23, and 29–30. The minimum range sequence of *A* is defined as *ρ*(*A*) = [10, 13, 23, 23, 29, 30].

#### Definition 8

(Range Compressed Row Storage). *For a sparse matrix* **A** *and its sorted CRS representation* (*C, R*), *we define Range Compressed Row Storage or rCRS*(**A**) *as two 1D arrays* (*C*′, *R*′) *which are called* column indices *and* row map *arrays respectively:*

▂ *C*′ = *ρ*(*C*_0_) *· · · ρ*(*C*_*n−* 1_), *where C*_*i*_ *is block i in the row decomposition of C;*
▂ *R*′ *is an array of length n* + 1, *in which R*′(*i*) *specifies the start index of ρ*(*C*_*i*_). *The last value is defined as R*′(*n*) = |*C*′|.

#### Remark 9.

In the column indices array *C*′ of rCRS, each element can be identified as either a lower or an upper bound of a range based on its index. Specifically, odd indices refer to lower bounds, whereas even indices refer to upper bounds.

#### Remark 10.

For each row decomposition block 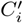 of the column indices array in rCRS, for *i* ∈ [0, *n*), the elements are sorted and the ranges indicated by consecutive pairs are disjoint. It can be shown that rCRS can be constructed from sorted CRS in linear time with respect to the number of non-zero values. Accessing element *a*_*ij*_ in **A** represented in rCRS reduces to determining whether *j* lies within any range in row *i*. Since the ranges are disjoint and sorted, this operation can be performed using binary search which requires 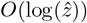 time in the worst case, where 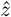 is the maximum size of 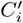 across all rows *i*. Note that, unlike CRS format, the size of *C*′ in rCRS is no longer equal to nnz(**A**). Appendix B provides detailed analysis of the rCRS format: Appendix B.1 explores the construction algorithm, while Appendix B.2 and Appendix B.3 analyze query time and space complexity, respectively.

The space complexity of the rCRS representation decisively depends on the distribution of non-zero values in the rows of the matrix. This representation can substantially compress the column indices array *C* – as low as *O*(*n*) compared to *O*(*nnz*(*A*)) in CRS – if the array contains long distinct ranges of consecutive integers. The limitation of this representation is that it can be twice as large when there is no stretch of consecutive indices in *C*. It is important to note that the compression rate can directly influence query time, as it is proportional to the compressed representation of non-zero values.

To mitigate this issue, we define a variant of rCRS, which is referred to as *Asymmetrical Range CRS* (*aCRS*). In the worst case scenario, the aCRS format takes as much space as the standard CRS while maintaining the same time complexity 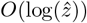 for query operations. We elaborate on the aCRS definition and related analysis in Appendix C.

Although aCRS theoretically provides a more compact representation compared to rCRS, the experimental results (in Section 5) show that it occupies as much space as rCRS when storing distance index 𝒯 for sequence graphs in practice. This is due to the fact that the ordering defined in Section 4.1, together with the topology of sequence graphs, rarely results in isolated column indices. Consequently, aCRS holds two integers per range, similar to rCRS. This behaviour can be observed in the index constructed for a chain graph (Figure 3), which often resembles the general structure of sequence graphs. Considering this fact and the relative ease of implementing efficient sparse matrix algorithms with rCRS compared to aCRS, particularly on GPU, we base our implementation on the rCRS format.

### 4.3 Range Sparse Boolean Matrix Multiplication (rSpGEMM)

Given sparse matrices **A**_*m×n*_ and **B**_*n×s*_ where *m, n*, and *s* are positive integers, SpGEMM is an algorithm that computes **C** = **A** *·* **B** using sparse representations of two input matrices. In this section, we introduce a new SpGEMM algorithm, called *Range SpGEMM* or *rSpGEMM* for short, computing matrix-matrix multiplication when input and output matrices are in the rCRS format. The algorithm is designed to scale well on both multi-core architectures (CPUs) as well as many-core architectures (GPUs).

Similar to most parallel SpGEMM algorithms, our rSpGEMM follows Gustavson’s algorithm [24]. Algorithm 2 illustrates its general structure, in which *A*(*i*, :) refers to row *i* of matrix **A**. This notation can be generalized to *B*(*j*, :) and *C*(*i*, :) in Algorithm 2 specifying row *j* and *i* in *B* and *C*, respectively. Note that Algorithm 2 only iterates over non-zero entries of **A** and **B**.

The algorithm computes **C** one row at a time. To compute *C*(*i*, :), it iterates over non-zero values in row *A*(*i*, :) and computes its contribution to *C*(*i*, :) by multiplying it to non-zero values in row *B*_*j*_; i.e. the term *a*_*ij*_*· B*(*j*, :) in Line 3. This contribution is then *accumulated* with the partial results computed from previous iterations. The row accumulation can be simplified to *C*(*i*, :) ← *C*(*i*, :) + *B*(*j*, :) in Boolean matrices as *a*_*ij*_ is 1.

Since computing each row of the final matrix is independent of the others, **C** = **A** *·* **B** can be seen as multiple independent vector-matrix multiplication *C*(*i*, :) = *A*(*i*, :) · **B**. For simplicity, we will only focus on the equivalent vector-matrix multiplication.

#### Algorithm 2

Gustavson’s algorithm.

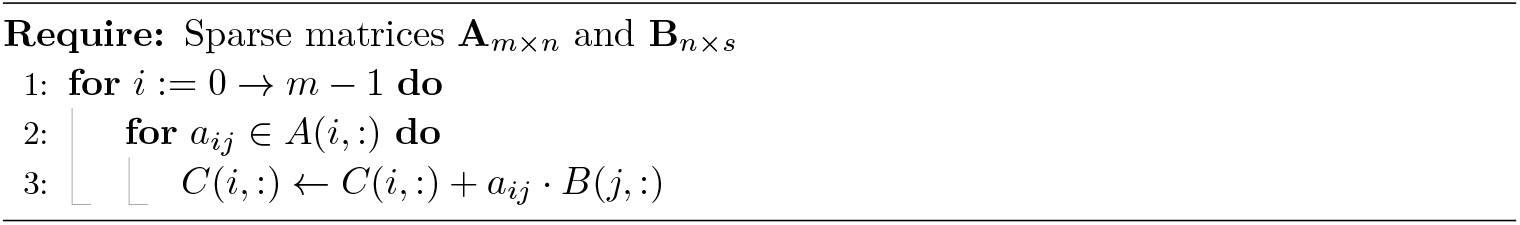

Given that the output matrix is also in the sparse format, knowing the number of non-zero values in each row of **C** is essential before storing the calculated *C*_*i*_ in the final matrix. This issue is usually tackled using a two-phase approach: firstly, the *symbolic* phase, delineates the structure of **C**, primarily corresponding to the computation of its row map array. Secondly, the *numeric* phase, carries out the actual computation of **C**.

Similar to SpGEMM algorithms based on Gustavson’s, the fundamental aspect of rSpGEMM resides in three anchor points: the data structure employed for row accumulation, memory access pattern to minimize data transfer latency, and distribution of work among threads in hierarchical parallelism. In the following, we address each of these points.

#### 4.3.1 Bi-Level Banded Bitvector as Accumulator

The row accumulation in Boolean SpGEMM, explained in Section 4.3, can be seen as the union of all column indices in *B*(*j*, :) for all *j* where *a*_*ij*_ ≠0. Figure 4a schematically depicts the row accumulation in computing a row of the final matrix. It is important to highlight that row accumulation in rSpGEMM is carried out on ranges of indices that are, by definition, disjoint and sorted (Remark 10).

**Figure 4.**
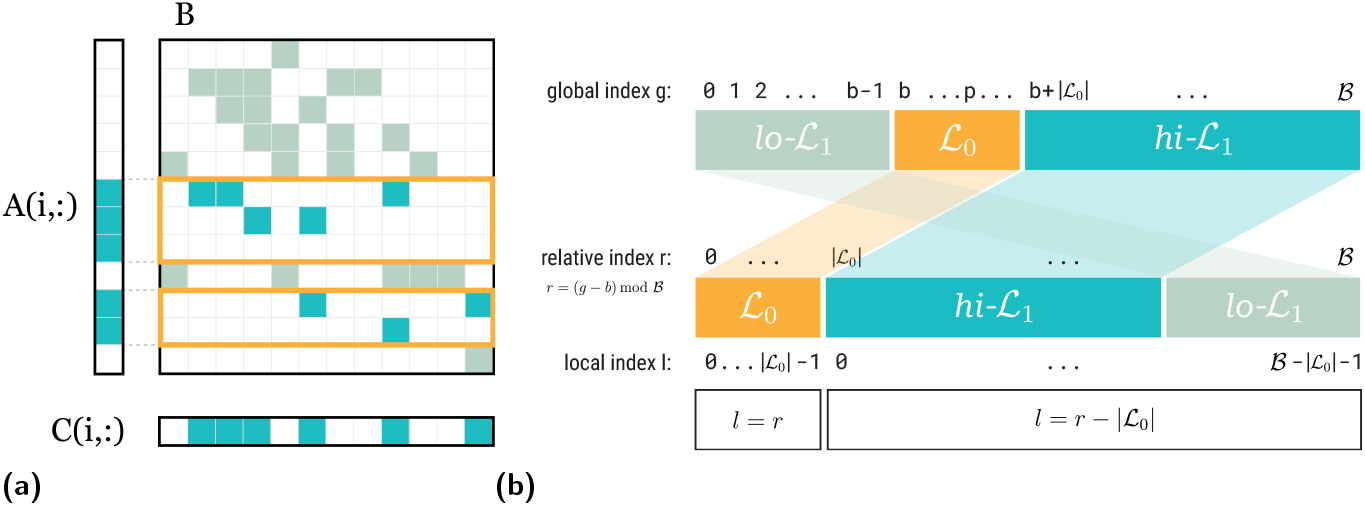
(a) Row accumulation in Boolean matrices can be viewed as the union of all non-zero values in rows of **B** that are specified by the non-zero values in *A*_*i*_ (rows highlighted by yellow rectangles). (b) BBB hierarchical structure and indexing: the assignment of column indices in a row to each level (top); bitvector rearranged according to *relative indices* (middle); the physical memory space corresponding to each level accessible with *local indices* (below).

Our method employs a dense bitvector, namely *Bi-level Banded Bitvector* (BBB), as an accumulator in rSpGEMM. From a high-level point of view, this bitvector supports two main operations: `scatter` and `gather`. The `scatter` operation stores a range of column indices at once, which essentially involves setting a range of sequential bits in the bitvector to one. Conversely, the `gather` operation retrieves the union of column indices stored by the `scatter` method in the form of the minimum range sequence (Definition 6). Computing *C*(*i*, :) is essentially a combination of `scattering` partial results and `gather`ing the final accumulated entries using BBB.

The `scatter` operation is a bit-parallel operation that sets bits corresponding to index range [*s, f*] in a series of *W*-bit operations. Each set bit in the bitvector indicates absence/presence of the corresponding column in the final result. The pseudocode for the `scatter` operation is given in Algorithm 3.

Each cluster of set bits in the final bitvector represents an interval in the final row *C*(*i*, :), with each interval corresponding to an entry pair in the rCRS format. In the symbolic phase, the `gather` operation counts the number of entries in rCRS with regards to non-zeros in row *C*(*i*, :). In the numeric phase, it constructs the resulting row as a minimum range sequence. Furthermore, in order to avoid scanning all words for set bits, the absolute minimum (*j*_min_) and maximum (*j*_max_) word indices are tracked and the final scan is limited to [*j*_min_, *j*_max_]. Since each row in **B** is sorted, *j*_min_ and *j*_max_ are updated once per row. The key advantage of the `gather` operation is that it relies solely on trivial bitwise functions, which – similar to `scatter` – exploit bit-level parallelism to convert clusters of set bits into their corresponding start and end indices. Algorithm 4 demonstrates the pseudocodes of the `gather` operations in the symbolic and numeric phrase.

The `gather` operation functions using four trivial bitwise functions: `cnt`, `map01`, `map10`, and `sel`. The `cnt` function simply counts the number of set bits in a word and is performed by a single machine instruction (`POPCNT`).

##### Algorithm 3

Bi-level Banded Bitvector `scatter` operation.

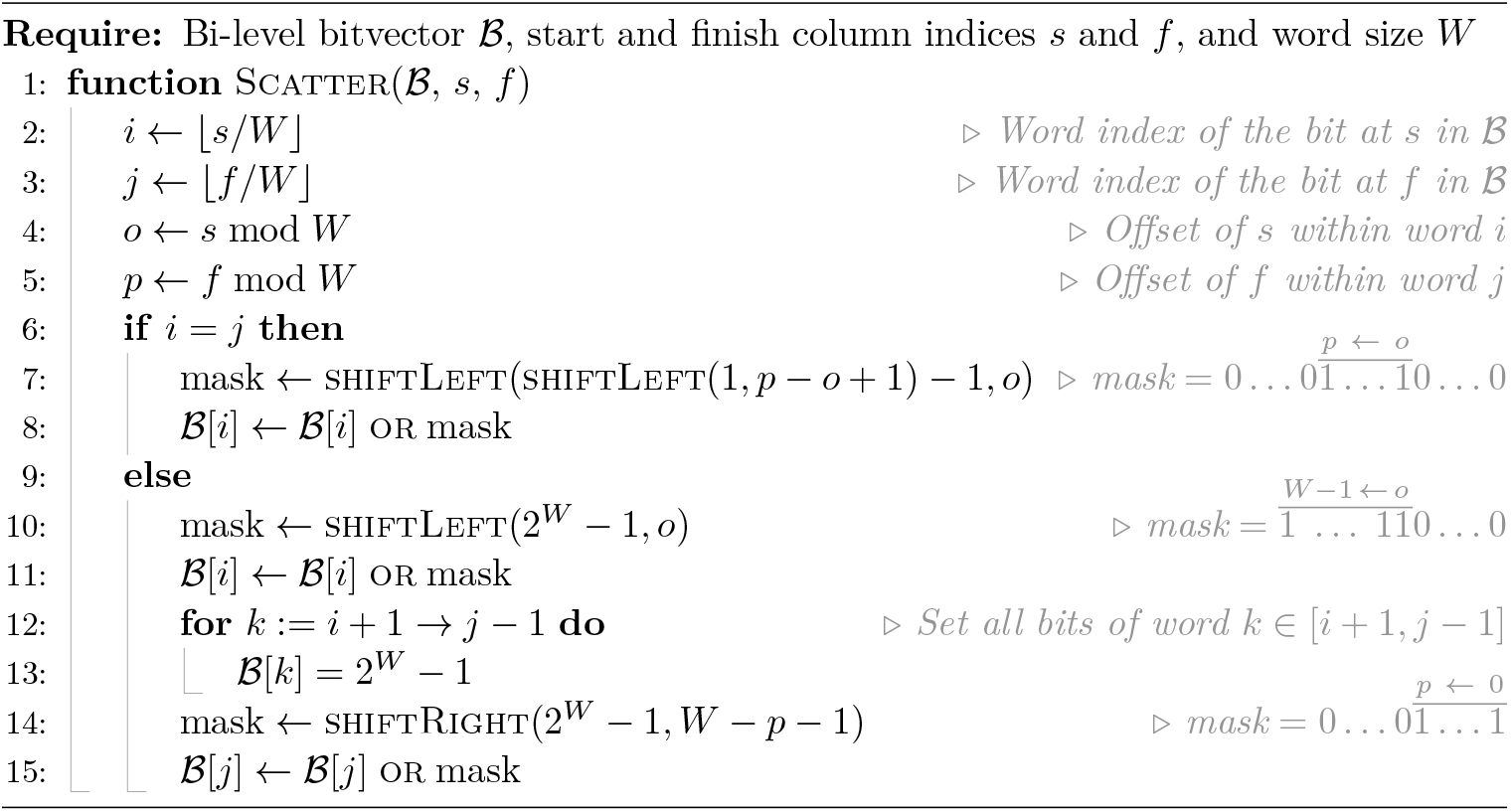

The `map01rr` and `map10` functions identify boundaries of consecutive set bits (runs of 1s) in a word through specific bit pattern transformations. The `map01rr` function detects 01 patterns, mapping them to 01 (i.e. itself) while mapping all other two-bit patterns (`00, 10`, and `11`) to `00`. Since a `01` pattern occurs when a 0 is followed by a 1, this function effectively marks the starting position of each run of consecutive 1s.

Conversely, the `map10` function detects `10` patterns, mapping them to `01` and all other patterns to `00`. Since a `10` pattern occurs when a 1 is followed by a 0, this function marks the position immediately after the end of each run of consecutive 1s. Both `map01rr` and map10 can be calculated using basic bitwise operations^1^ in constant time independent of the word size. The border cases, when `01` or `10` occur at the boundaries between two words, are handled by peeking at the immediate bit on the left of the bit that is being processed. In other words, the most significant bit (MSB) of the previous word is appended to the current word before applying `map*` functions. This bit, which is called *carried* bit, is zero for the first word.

Finally, the `sel` (*x, i*) function gives the position of *i*-th set bit in word *x* relying on native machine instructions (e.g. `fns` intrinsic in CUDA and `PDEP` and `TZCNT` instructions on CPUs). The shiftLeft(*w, l*) and shiftRight(*w, l*) function calls in Algorithm 3 and Algorithm 4 denote bitwise left-shift and right-shift operations of word *w* by *l* bits, respectively. In the symbolic phase, `gather` counts the number of entries in rCRS with regards to non-zeros in row *C*(*i*, :). This count is equivalent to twice the number of 01 occurrences in the bitvector and can be formulated as `cnt` (`map01rr` (*w*)) for all modified words *w* in the bitvector. In order to convert the set bit stretches to integer intervals in numeric phase, the first and the last bit of each stretch are marked by applying `map01rr` and `map10` functions on all words. Finally, calling `sel` on each set bits in (`map01` (*w*) or `map10` (*w*)) gets the final minimum range sequence of the accumulated indices.

##### Algorithm 4

Bi-level Banded Bitvector `gather` operation.

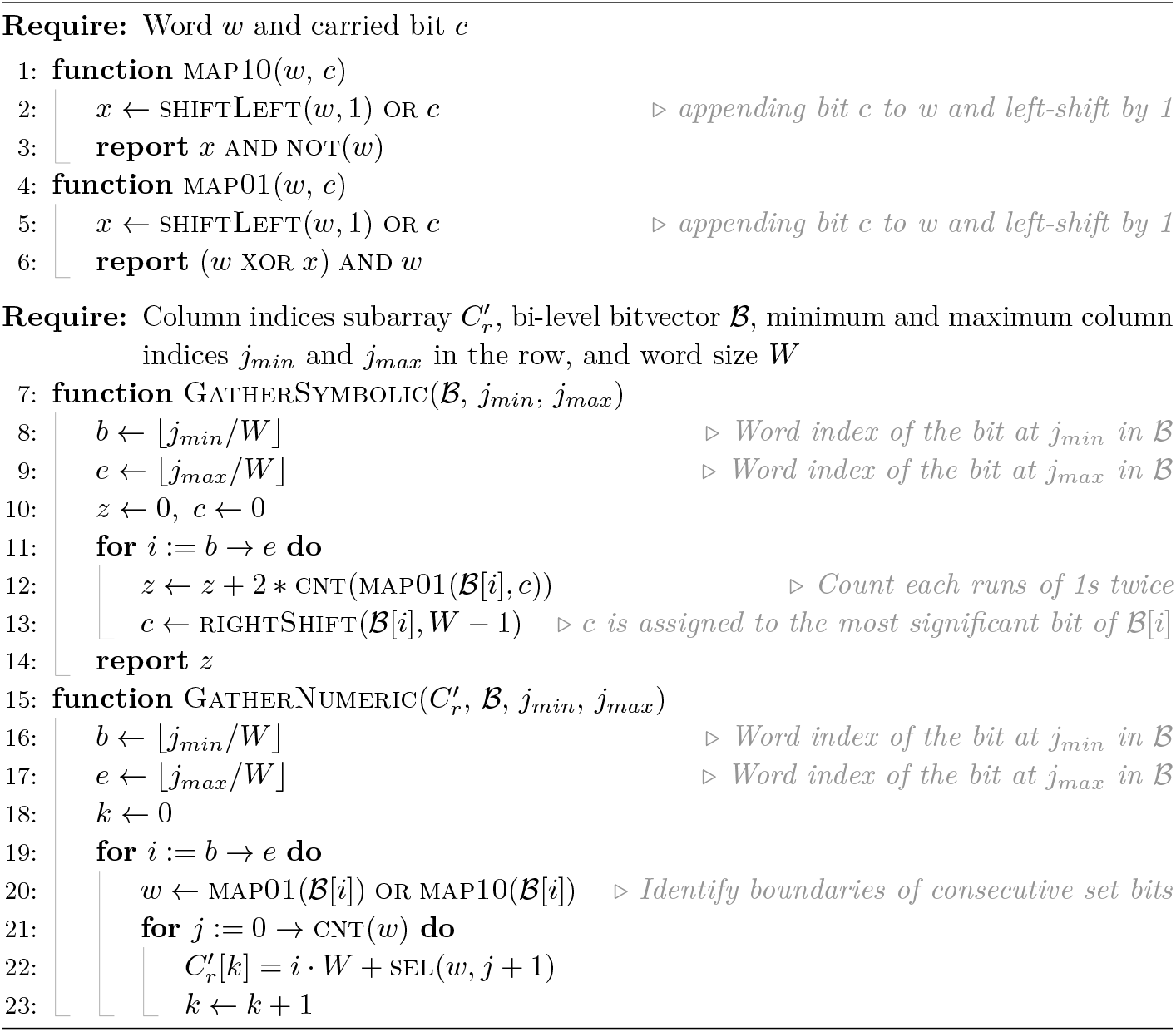

##### Example 11.

Consider the 16-bit word *w* = `0011110011001110` representing the bitvector resulted from `scatter` operations, which essentially denotes the absence/presence of column indices in the final row. Then, we have `map01rr` (*w*) = `0010000010001000`, in which all bits except the occurrences of 01 are set to 0 and `map10` (*w*) = `0000001000100001`, in which all occurrences of 10 are retained as 01 and the rest of bits are set to zero. It can be seen that the number of intervals with consecutive set bits in *w* (highlighted by underline) is equal to `cnt` (`map01rr` (*w*)) = 3. Calling `sel` on all set bits in (`map01` (*w*) or `map10` (*w*) = `0010001010101001`) yields the minimum range sequence of the column indices present in the bit vector; i.e. [2, 5, 8, 9, 12, 14]. Note that the upper bounds are substracted by one to represent closed intervals.

##### Non-blocking Hierarchical Parallelism

All `scatter` operations for each pair of [*s, f*] are computed in parallel across multiple threads. Concurrent write accesses to the bitvector can lead to race conditions depending on how computations are *partitioned* among threads. Different *partitioning schemes* might impose different requirements which will be explained in Section 4.3.2. In a particular partitioning scheme, if a race condition is possible, atomic operations are opted for all or a subset of parallel accesses.

Similarly, the `gather` operations are performed in parallel across all words. This parallelization introduces a synchronization challenge: each thread needs to know the starting index in the column indices array where the minimum range sequence of the assigned word should be stored, but this index depends on the range sequence sizes of all preceding words. Rather than introducing thread synchronization overhead, each thread independently re-computes its required starting index.

Apart from distributing `scatter` or `gather` operations among threads, BBB can benefit from another level of parallelism: vector parallelism, or single-instruction-multiple-data (SIMD), if supported by the underlying architecture. Vector parallelism allows increasing the effective word size by conducting `scatter` or `gather` operations simultaneously on multiple machine words. The length of SIMD instructions, i.e. the *vector length*, is architecture-dependent and usually matches the size of the cache line.

##### Hierarchical Memory

The bitvector is designed to perform well in scenarios where non-zeros in row *C*(*i*, :) are localized, which reflects a practically common scenario. To this end, two design decisions have been made. Firstly, the size of the bitvector is bounded by the bandwidth observed in *C*(*i*, :) which is calculated before the symbolic phase. Secondly, the bitvector is partitioned into two levels ℒ_0_ and ℒ_1_ in order to minimize the memory latency by utilizing the locality of non-zero values in *C*(*i*, :). The first level (ℒ_0_) is allocated on the fast (low-latency) memory if there is hardware support (e.g. shared memory in GPUs). The rest of the bit fields is mapped to the second level ℒ_1_ and allocated on the larger memory but with higher latency (e.g. global memory on GPU). On hardware where software-managed memory hierarchy is not available (like CPUs), this model promotes cache-friendly memory accesses.

The rationale behind this design is that the first level ℒ_0_ spans the range of bits that are more likely to be accessed during row accumulations. If all non-zero values in a row are bound to ℒ_0_ index range, no words in ℒ_1_ are fetched or modified. Therefore, all `scatter` and `gather` operations mentioned earlier are performed on words in the fastest memory. The relative position of the first level within a row is specified by the index of its center bit *p*, referred as *pivot bit*. For example, the center of the band in each row of *C* is a viable option for *p*. In this work, we chose *p* = *i* to set the ℒ_0_ region around the diagonal as node *i* is more likely to be adjacent to nodes with ranks close to *i* in adjacency matrices.

The arrangement of bits in two levels necessitates calculating internal (local) indices within levels from column (global) indices. Local indices in each level are calculated from corresponding global indices relative to *p* and using modular arithmetic. Figure 4b demonstrates the assignment of column indices in a row to each level (top), bitvector rearranged according to *relative indices* (middle), and the physical memory space corresponding to each level accessible with *local indices* (below).

#### 4.3.2 Partitioning Schemes

Consider computing **C** = **A***·* **B** using rSpGEMM with matrices in rCRS format, there are different ways to distribute required work units among available threads. The partitioning and assignment of work units to computational units can influence the runtime, memory usage, and memory access patterns. Additionally, our implementation benefits from the Kokkos library [13] to achieve performance portability, meaning that it can be executed on multiple supported hardware architectures. As different hardware can impose different requirements, an efficient partitioning strategy on a specific architecture is not necessarily efficient on another.

Despite the inherent differences in their design, computational hardware can be abstracted by a unified logical view. This abstraction allows us to effectively describe the partitioning schemes and the hierarchical parallelism inherent in multi-core and many-core architectures. Processing units typically comprise multiple or many *cores*, each of which is designed to execute multiple threads concurrently. In this unified computational model, a group of threads executing within a single core is termed a *team of threads* and the number of supported threads is called *team size*. Within a team, threads might share memory resources and be able to synchronize. Moreover, each thread might be capable of executing vector operations.

Vector parallelism leverages the simultaneous processing of multiple data entries within a single instruction cycle (SIMD or vectorization on CPU and *warps* in GPU). The number of vector lanes that a thread can handle is defined as the *vector size*. We also define *block size* as the total parallel processing capability of a core; that is, *team size* × *vector size*.

Inspired by [13], we examined different strategies: *Thread-Sequential, Team-Sequential*, and *Thread-Parallel* partitioning schemes:

##### Thread-Sequential

In this partitioning scheme, each team is responsible for computing a range of rows in **C**, with these rows distributed across threads within the team. To compute *C*(*i*, :), a single thread *sequentially* iterates over ranges in *A*(*i*, :). For each range [*j, k*] in *A*(*i*, :), the thread accumulates all column indices corresponding to non-zero elements in rows *B*(*j*, :) through *B*(*k*, :). The thread exploits vector parallelism for storing each range in the accumulator.

Figure 5 illustrates the distribution of work units across threads in this scheme. In this example, each team contains two threads with vector size 2, and two rows are assigned to each team. The figure shows the access patterns of Threads 1 and 2 to **B**, where they sequentially traverse rows corresponding to non-zero values in their assigned row of **A** (indicated by unique symbols) and accumulate non-zero elements in these rows. The remaining access patterns are omitted for clarity.

**Figure 5.**
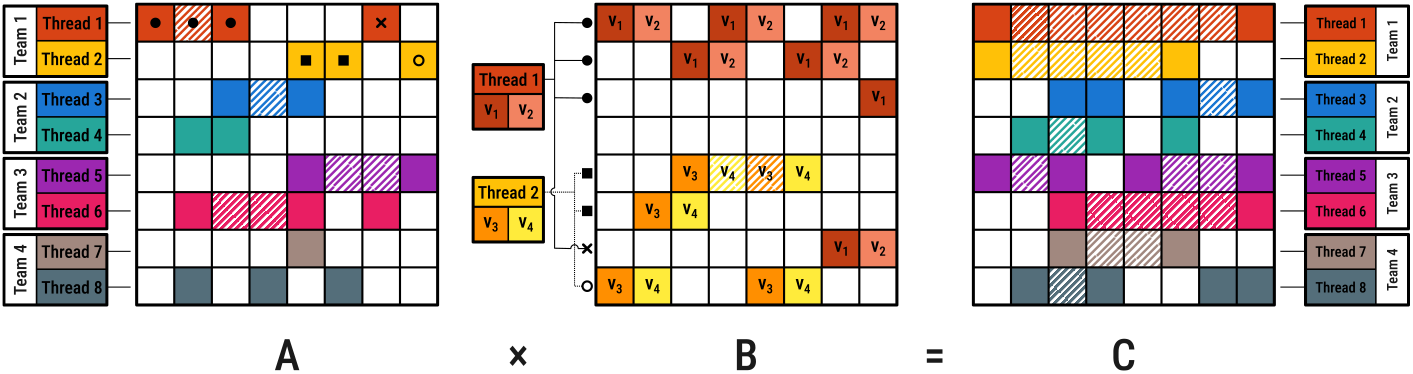
Thread-Sequential partitioning: work distribution and memory access patterns. Threads sequentially accumulate non-zero elements in rows of **B** corresponding to ranges in their assigned rows of **A**, indicated by unique symbols (filled squares, circles, crosses, and empty circles), with only Threads 1 and 2 shown for clarity.

Since all threads work with separate accumulators, no data race conditions occur. Hence, no synchronization is needed. However, this approach requires more memory as the number of active threads increases. This may be restrictive on GPUs for larger matrices due to limited global memory on devices.

##### Team-Sequential

As shown in Figure 6, each team is assigned to a single row of **C** in this scheme. As suggested by the name, each team *sequentially* iterates over ranges in *A*(*i*, :) as well as the corresponding rows in **B**. All threads in a team are assigned to different ranges within a single row of **B**. Similar to Thread-Sequential, `scatter` and `gather` operations on bitvectors utilize vector parallelism.

**Figure 6.**
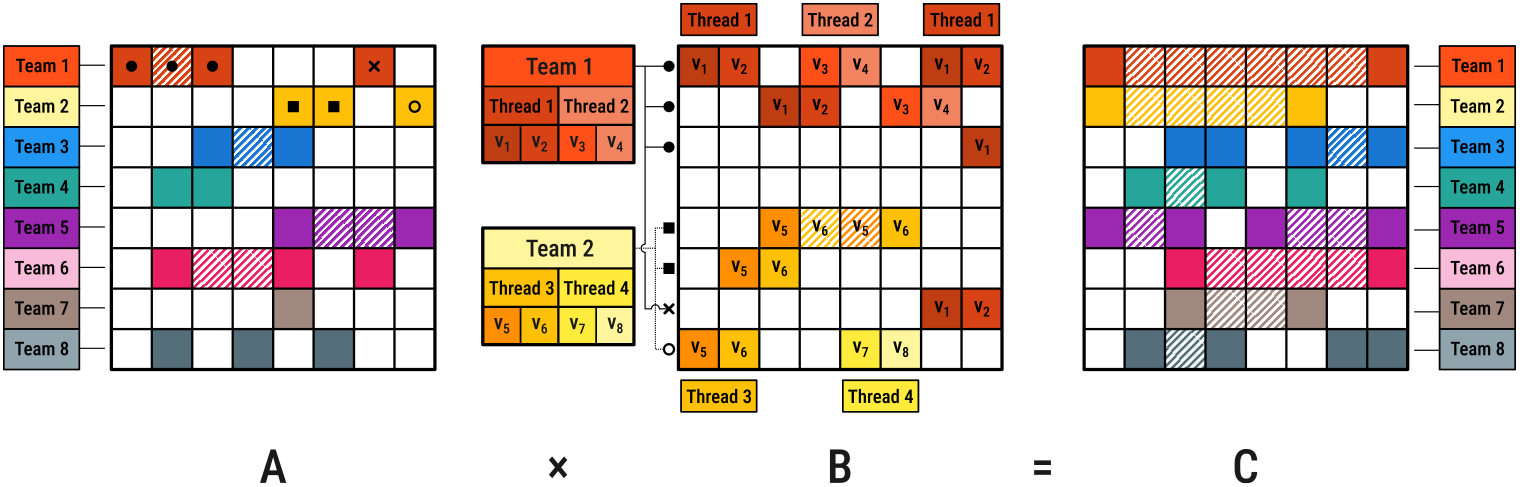
Team-Sequential partitioning: work distribution and memory access patterns. Teams are assigned to individual rows of **C**, with threads within each team processing different ranges within a single row of **B**, with only Team 1 and 2 shown for clarity.

Since all team threads work on a row in **B**, there are fewer active rows compared to Thread-Sequential. Therefore, it has a smaller memory footprint. However, when **B** is very sparse, this scheme may not fully utilize available computational resources if there are fewer ranges per row than threads in the team.

Moreover, as ranges in each row of **B** are disjoint, there is no race condition in row accumulation except for the first and last words of each range. Therefore, bitwise operations on those words must be atomic.

##### Thread-Parallel

Similar to Team-Sequential, each team is assigned to compute a single row of **C** in this scheme, with each team sequentially traversing ranges in *A*(*i*, :). However, for a range [*j, k*] in *A*(*i*, :), all ranges in rows from *B*(*j*, :) to *B*(*k*, :) are distributed among threads. As a result, underutilization is less likely in this approach as long as the number of non-zero values (not ranges) is sufficiently large in **A**. As with previous approaches, threads use vector parallelism when computing each ranges in **B**. A schematic view of this partitioning is shown in Figure 7.

**Figure 7.**
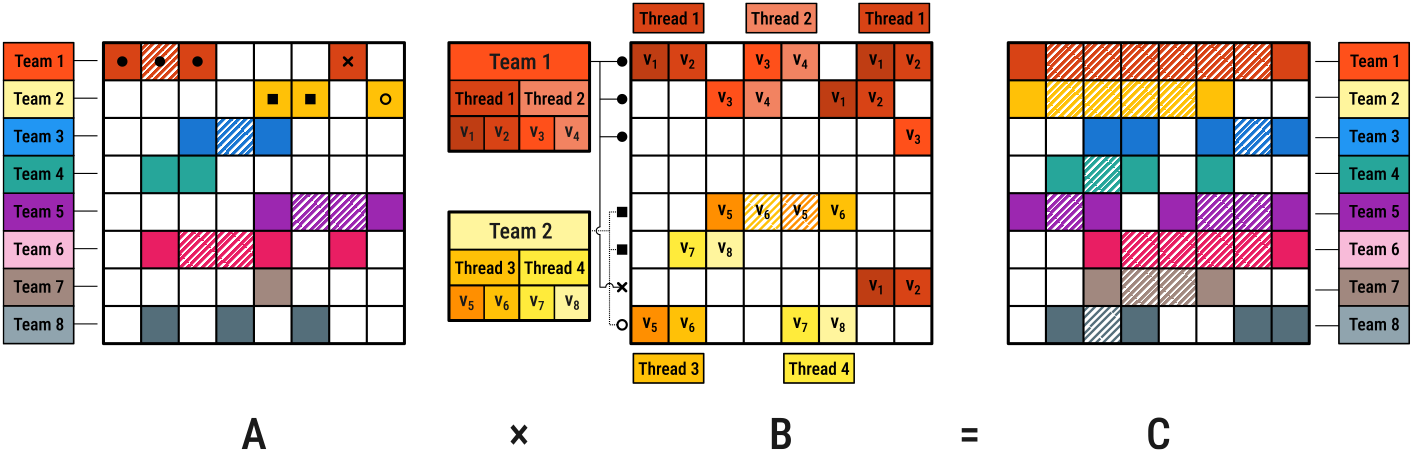
Thread-Parallel: work distribution and memory access patterns. Each team is assigned to a single row of **C**, with threads within the team processing multiple rows of **B** corresponding to a single non-zero range in the assigned row of **A**, indicated by unique symbols (filled squares, circles, crosses, and empty circles), with only Team 1 and 2 shown for clarity.

In terms of memory requirements, this scheme is similar to Team-Sequential. However, unlike Team-Sequential where threads work within a single row, ranges from multiple rows can overlap, requiring all bitwise operations for row accumulation to be atomic to prevent race conditions.

### 4.4 Range Sparse Boolean Matrix Addition (rSpAdd)

In this section, we shift our focus to calculating matrix-matrix addition in rCRS format which is required for computing the distance index. We propose the *Range Boolean Sparse Add* (or *rSpAdd*) algorithm which computes matrix **C** = **A** ∨ **B** where all matrices are in rCRS format and ∨ is Boolean matrix addition defined in Definition 2.

Considering that non-zero values in each row of rCRS matrices are represented by ranges, each with two integers (Remark 9), and these integers are sorted and disjoint (Remark 10), row *C*(*i*, :) is equivalent to the sorted combination of ranges in both rows *A*(*i*, :) and *B*(*i*, :), where overlapping ranges are collapsed into one. For each row in **C**, this is achieved by using two pointers, each initially pointing to the first element of the respective rows. The algorithm peeks at the pairs in *A*(*i*, :) and *B*(*i*, :) indicated by the pointers. If two ranges are disjoint, it inserts the one with smaller bounds in *C*(*i*, :) and increments the pointer indicating the inserted pair by two. If two ranges overlap, they will be merged by inserting the minimum of lower bounds and the maximum of upper bounds of the two pointed elements into *C*(*i*, :) in that order. Then, both pointers are incremented to indicate the next pairs. In case either pointer reaches the end of the row, the remaining elements from the other row are directly appended to *C*(*i*, :).

The time complexity of this algorithm is bound by compressed representation of input matrices in rCRS format. Since the ranges in rCRS are disjoint and sorted, merging two rows *A*(*i*, :) and *B*(*i*, :) can be done in Θ(|*A*(*i*, :) | + |*B*(*i*, :) |); in which |*A*(*i*, :) | and |*B*(*i*, :) | are the sizes of *A*(*i*, :) and *B*(*i*, :) in rCRS format, respectively. rSpAdd not only requires smaller working space compared to SpAdd but is arguably faster due to compressed representation of the matrices.

### 4.5 Distance Index

As stated in Section 3.1, the distance index constructed by Equation (1) is a block-diagonal matrix. For this reason, DiVerG builds the distance index incrementally for each block; i.e. computing the distance matrix for each component of the sequence graph individually. The adjacency matrix for each component can be constructed in rCRS format one row at a time. Therefore, it is not required to store the whole matrix in sorted CRS format to be able to convert it to rCRS. Once matrices **A** and **I** are in rCRS format, 𝒯 can be computed using rSpAdd and rSpGEMM. The powers of **A** and **A** ∨ **I** are computed using “exponentiation by squaring” [29]. Ultimately, matrix 𝒯 is further compressed by encoding both the column indices and row map arrays using Elias-*δ* encoding [17].

DiVerG provides an exact solution to the DVP through its compressed format for storing both final and intermediate matrices when computing the distance index 𝒯. The compression ratio directly impacts the efficiency of the matrix multiplication operations, with higher compression resulting in faster construction. Since compression efficiency depends on the topology of the sequence graph and the structure of its adjacency matrix, establishing tight theoretical bounds on space and time complexities is challenging. However, we can derive meaningful lower bounds to facilitate comparison with previous approaches.

It can be demonstrated that for a sequence graph *G* = (*V, E*), indexing cost is lower bounded by that of the chain graph *G*_*c*_ = (*V* ′, *E*′) representing the longest path in *G* [28]. This analysis applies to our rSpGEMM algorithm as well. For a chain graph, the index 𝒯 has a particularly simple structure (Figure 3): in each row *i*, all columns from *d*_1_ to *d*_2_ is 1. This structure offers two advantages in the rCRS format:

#### Space complexity

Each row requires only two integers regardless of the distance range, resulting in Ω(|*V*|) space complexity – independent of distance parameters *d*_1_ and *d*_2_, unlike PairG’s Ω(|*V*| *· d*) where *d* = *d*_2_ *− d*_1_. Note that Elias encoding of the row map and column indices arrays compresses the final index even further in practice.

#### Time complexity

While rSpGEMM performs the same operations as SpGEMM, bit-parallelism reduces the computation time by a factor of the word size *W*. Jain et al. [28] showed that computing 𝒯 for *G* using SpGEMM requires Ω (|*V* ′ | *d*^2^ + log *d*_1_)); therefore, DiVerG’s rSpGEMM requires Ω (|*V* ′| (*d*^2^*/W* + log *d*_1_)). Note that, this improvement is particularly significant when computing 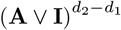, where multiple non-zero values per row enable effective bit-parallelism. Additionally, query operations are improved from *O*(*d*) to *O*(1) complexity, where *d* = *d*_2_ *− d*_1_, as each row contains exactly two integers. Given that sequence graphs often closely resemble their linear structure, these bounds provide a reasonable approximation of actual performance while highlighting the fundamental advantages of our approach.

#### 4.5.1 Multi-step indexing and merge operation

The rationale behind preprocessing the genome graph is to construct an index once and use it to efficiently answer queries multiple times. It is important to note that the fragment model may vary between samples. Therefore, the index may need to be adjusted accordingly. To avoid re-indexing large genomes, the distance index can be constructed for a set of disjoint distance constraints. These chunks can be used as building blocks and can be efficiently merged to construct a new index with different constraints.

##### Lemma 12

(Distance Index Merge). *Given two distance indices* 𝒯_1_ *and* 𝒯_2_ *for distance constraints* [*d*_1_, *d*_2_] *and* [*d*_3_, *d*_4_] *respectively, the merged matrix* 𝒯_1_ ∨ 𝒯_2_ *is a distance matrix for constraints* [*d*_1_, *d*_2_] ∪ [*d*_3_, *d*_4_], *where* ∨ *is Boolean matrix addition*.

If two indices are represented in rCRS format, this operation takes *O*(rnnz(𝒯_1_) + rnnz(𝒯_2_)) time, where rnnz(𝒯_*i*_) is the number of entries of 𝒯_*i*_ in rCRS format. This computation is often faster than directly reconstructing the index for the union of the distance constraints.

## 5 Experimental Results

### 5.1 Implementation

DiVerG is implemented in C**++**17. While we develop our theory using character graphs, our implementation handles general string-labelled sequence graphs without loss of generality. The implementation stores all matrices – including the adjacency matrix for the input graph, intermediate matrices, and the final distance matrix 𝒯 – in rCRS data structure. The matrix 𝒯, in the final stage, is converted to *encoded rCRS* format (described in Section 4.5) whose size is reported as *index size* in our experiments (Section 5.4). Our implementation relies on Kokkos [13] for performance portability across different architectures, and we specifically focused on two execution spaces: OpenMP (CPU), and CUDA (GPU). Despite the fact that algorithms implemented in the Kokkos programming model are portable across various platforms, specific optimizations and design decisions are tailored for particular architectures. We compared the performance of our algorithm with PairG which uses the `kkSpGEMM` meta-algorithm implemented in Kokkos [13] for computing sparse matrix multiplication.

Kokkos offers several algorithms for computing SpGEMM. The `kkSpGEMM` algorithm attempts to choose the best algorithm based on the inputs. As the PairG implementation was in practice not runnable on GPU, we re-implemented the corresponding part of PairG to support GPU and used this implementation for comparison.

All CPU experiments were conducted on an instance on de.NBI cloud with a 28-core (one thread per core) Intel® Xeon® Processor running Ubuntu 22.4. Distance index construction on GPU were carried out on an instance with an NVIDIA A40 GPU running Ubuntu 22.4.

### 5.2 Tuning Meta-parameters

We implemented the rSpGEMM algorithm with different partitioning schemes and evaluated the performance of each using various block sizes, as described in Section 4.3.2. In this work, we mainly report the performance of DiVerG with the best partitioning scheme and block size for each architecture. Table 1 summarizes the selected partitioning schemes and block sizes for each execution space and input data size. The implementation automatically determines these meta-parameters based on the underlying platform (CPU or GPU) and the input graph. An input graph is considered *large* if its total sequence length is larger than 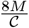, where *M* is the size of global memory on GPU in bytes and is the number of concurrent blocks that can run on GPU (or the number of CUDA cores). This threshold is based on a rough estimation of the total memory required for all active row accumulators as well as working space for storing intermediate matrices. Further tuning is possible in individual cases depending on the hardware and the inputs.

**Table 1.** Selected partitioning and block sizes for different execution spaces and input sizes.

| Execution space (input size) | Block size |  |  | Partitioning |
| --- | --- | --- | --- | --- |
|  | team size | vector size | thread work size |  |
| OpenMP (all graphs) | 1 | 1 | - | Thread-Sequential |
| CUDA (large graphs) | 8 | 8 | - | Team-Sequential |
| CUDA (small graphs) | 8 | 8 | 8 | Thread-Sequential |

As explained in Section 4.3.2, the Thread-Sequential scheme does not scale well as it imposes substantial memory requirements for large graphs. However, by reducing the number of active rows through carefully choosing the block size and thread work size, the Thread-Sequential partitioning can perform as efficiently as Team-Sequential in most cases. As a guideline for block size selection, we suggest team size 32 and vector size 32, with thread work size *ω* calculated as 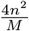 and rounded to the nearest power of 2, where *M* is the size of global memory on the GPU in bytes and *n* is the total sequence length of the graph.

### 5.3 Datasets

DiVerG is assessed using seven different graphs. Four of these graphs were previously employed by PairG in their evaluation, while three additional graphs have been introduced to address significantly larger and more complex scenarios. Together, these graphs span over a spectrum of scales, complexities, and construction methods.

Three of these graphs are variation graphs of different regions of the human genome, constructed from small variants in the 1000 Genomes Project [1] based on GRCh37 reference assembly: the mitochondrial DNA (mtDNA) graph, the BRCA1 gene graph (BRC), and the killer cell immunoglobulin-like receptors (LRC_KIR) graph. One graph is a de Bruijn graph constructed using whole-genome sequences of 20 strains of B. *anthracis*. We added three new variation graphs to this ensemble: the MHC region graph constructed from alternate loci released with GRCh38 reference assembly [37], and the HPRC CHM13 graph [31] constructed by minigraph. Table 2 presents some statistics of the sequence graphs used in our evaluation. More details about the datasets can be found in Appendix D.

**Table 2.** Statistics of the sequence graphs used for performance evaluation.

| Graphs | Nodes<sup>1)</sup> | Edges |
| --- | --- | --- |
| mtDNA (GRCh37-1KGP) | 21,037 | 26,842 |
| BRCA1 (GRCh37-1KGP) | 83,267 | 85,422 |
| LRC_KIR (GRCh37-1KGP) | 1,108,768 | 1,154,049 |
| MHC-minigraph (GRCh38-alt) | 10,987,753 | 11,078,552 |
| *B. anthracis* (20 strains) | 11,237,088 | 11,253,454 |
| HPRC CHM13 chr22 | 63,520,037 | 63,527,923 |
| HPRC CHM13 chr1 | 261,344,589 | 261,360,990 |

1) Note that all graphs are *character graphs*.

### 5.4 Results

We evaluate the *index size* (`size`), *construction time* (`ctime`), and high water mark memory footprint during index construction (`mem`) for different distance constraints using the graphs explained in Section 5.3. Moreover, the *query time* (`qtime`) is measured for the distance index by averaging the time spend for querying 1M randomly sampled paired positions throughout each graph. Our method is evaluated using two separate sets of distance constraints.

#### Performance Evaluation with Increasing Distance Constraints Intervals

In the first experiment, performance metrics are computed for two graphs, BRCA1 (small) and MHC (medium-sized), with distance constraints: (0, 128), (0, 256), (0, 512), and (0, 1024). This experiment reveals the behaviour of our algorithm as the distance range increases and specifically highlights the performance of rSpGEMM in computing the term 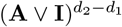 in Equation (1), which is a bottleneck in calculating 𝒯. The performance measures are computed on both CPU (OpenMP) and GPU (CUDA).

Figure 8a illustrates the performance measures and their comparison with PairG for the BRCA1 graph. As can be seen, DiVerG is about 4–7x faster in query time and 45–172x smaller in size in comparison to standard sparse matrix format and operations. Furthermore, the results show that DiVerG is approximately 9–28x faster than PairG in construction time on GPU and 10–68x on CPU. Moreover, the query time and index size in DiVerG do not change as the range of the distance constraints increases. This is because the number of non-zero values, i.e. ranges, in the rCRS format saturates quickly and stays constant.

**Figure 8.**
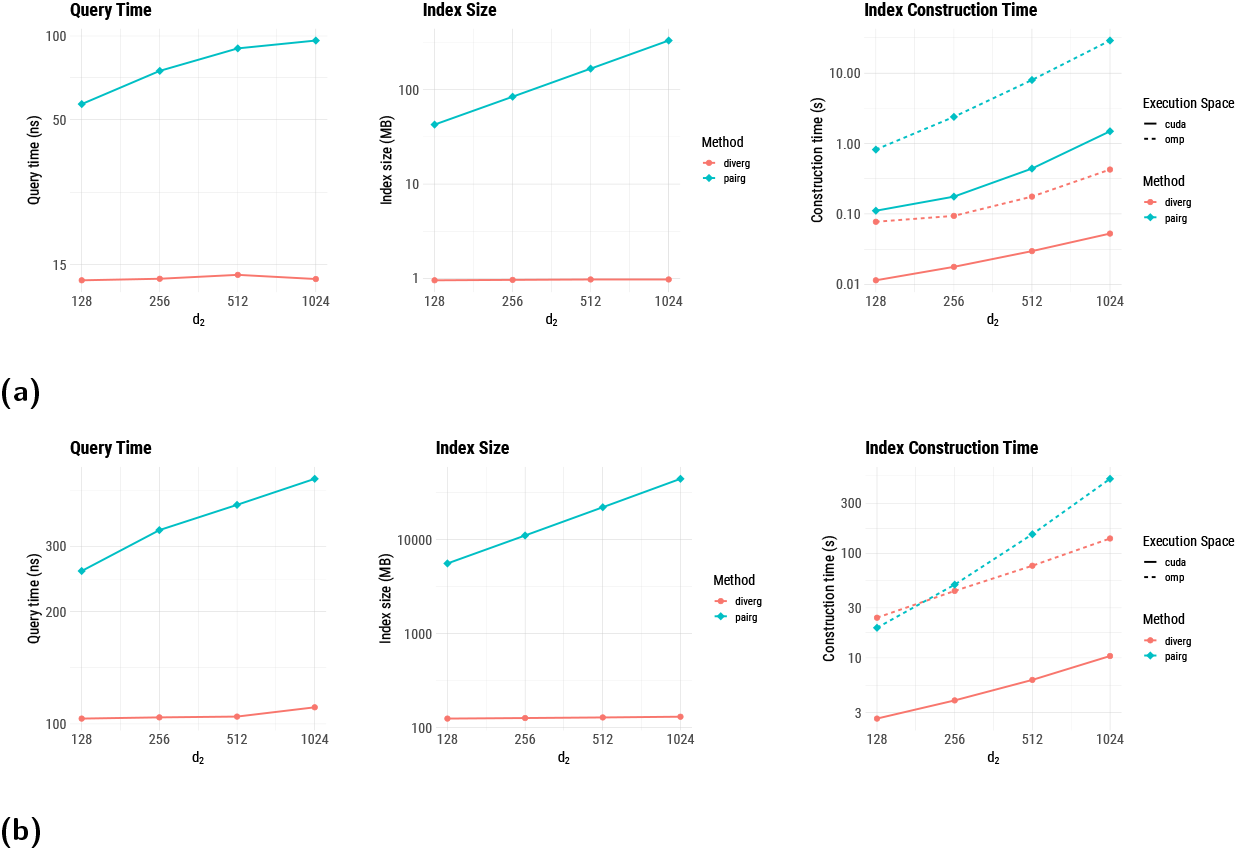
Log-scale performance of DiVerG for BRCA1 (a) and MHC graphs (b).

Our evaluations for the MHC region graph conducted with the same distance intervals are illustrated in Figure 8b. Although the MHC graph is much larger than the BRCA1 graph, the trends in the results remain consistent. Specifically, DiVerG achieves about 2.5–4x speed-up in accessing the matrix; the index in the rCRS format is approximately 44–340x smaller compared to the CRS format; and the query time and index size do not change with an increase in the distance interval. As PairG fails to construct the index on GPU due to the graph size and limited memory available on the device, we only report the runtime on CPU for this graph.

As shown in Figure 8b, DiVerG is faster that PairG in indexing the graph on CPU, except for the distance interval (0, 128). However, as the distance intervals grow larger, DiVerG becomes up to 3x faster in index construction. This speedup increases to 7–49.5x when DiVerG is executed on GPU: 7x for the distance constraint (0, 128) and 49.5x for (0, 1024).

#### Performance Evaluation with Offset Distance Constraints Intervals

In the second experiment, DiVerG is assessed for more realistic distance constraints (0, 250), (150, 450), and (350, 650), resembling the inner distance model for paired-end reads with fragment sizes 300 bp, 500 bp, and 700 bp, respectively. The evaluation is conducted on all graphs described in Section 5.3, including chr1 and chr22 of the HPRC CHM13 graph to demonstrate the scalability of DiVerG to large genomes. All benchmarks are executed on an NVIDIA A40 GPU which has 48 GB of global memory. For certain datasets and parameter configurations, PairG failed to construct the index, either due to excessive memory requirements or because it did not complete within a 6-hour time frame. In those cases, we only report the performance of DiVerG, and the size of the PairG index which is computed via direct conversion from rCRS to CRS. Given the size of the graphs in Table 2 and the number of non-zero entries in the final matrices, the size of the CRS format is computed by considering 4 bytes per column index, 4 bytes per value, and 8 bytes per entry in the row map array.

Table 3 shows the superior performance of DiVerG across all datasets and parameter settings, with indices that are about 170–420x smaller and, in instances where PairG successfully completes, created 3–6x faster with considerably lower memory requirements than PairG. The average number of column indices in each row in the rCRS format is about 2.1 for all variation graphs and all distance intervals. This number is approximately 4.8, 5.8, and 8.2 for the de Bruijn graph with distance intervals (0, 250), (150, 450), and (350, 650), respectively, while the average number of non-zero values per row in the equivalent CRS format is close to 300, i.e. *d*_2_*− d*_1_. This explains the high compression ratio of the DiVerG distance index. Additionally, query time shows a 3–5x speedup across all pangenome graphs. Our method can handle significantly larger graphs that are beyond the capabilities of PairG as it only requires a few tens of megabytes *auxiliary space* in memory per active thread.

**Table 3.** Performance of DiVerG compared to PairG on an NVIDIA A40 GPU in terms of index size (size), construction time (ctime), high water mark memory footprint (mem), and query time (qtime), along with the number of non-zero values in the final index (nnz).

| | $d_1-d_2$ | nnz | DiVerG | | | | PairG | | | |
| --- | --- | --- | --- | --- | --- | --- | --- | --- | --- | --- |
|  |  |  | size | ctime | mem | qtime | size | ctime | mem | qtime |
| mtDNA | 0–250 | 7M | **287KB** | **37<sub>ms</sub>** | **1MB** | **12<sub>ns</sub>** | 52MB | 215ms | 2.17GB | 36<sub>ns</sub> |
|  | 150–450 | 8M | **245KB** | **39<sub>ms</sub>** | **1MB** | **11<sub>ns</sub>** | 60MB | 240ms | 2.18GB | 37<sub>ns</sub> |
|  | 350–650 | 8M | **243KB** | **39<sub>ms</sub>** | **1MB** | **11<sub>ns</sub>** | 58MB | 239ms | 2.18GB | 36<sub>ns</sub> |
| BRCA1 | 0–250 | 21M | **1MB** | **35<sub>ms</sub>** | **4MB** | **13<sub>ns</sub>** | 164MB | 113ms | 542MB | 69<sub>ns</sub> |
|  | 150–450 | 26M | **1MB** | **41<sub>ms</sub>** | **5MB** | **13<sub>ns</sub>** | 197MB | 173ms | 500MB | 75<sub>ns</sub> |
|  | 350–650 | 26M | **1MB** | **44<sub>ms</sub>** | **5MB** | **13<sub>ns</sub>** | 197MB | 176ms | 500MB | 75<sub>ns</sub> |
| LRC | 0–250 | 290M | **13MB** | **320<sub>ms</sub>** | **52MB** | **25<sub>ns</sub>** | 2.2GB | 1.1s | 4.5GB | 113<sub>ns</sub> |
|  | 150–450 | 349M | **13MB** | **374<sub>ms</sub>** | **70MB** | **25<sub>ns</sub>** | 2.7GB | 1.9s | 5.4GB | 120<sub>ns</sub> |
|  | 350–650 | 350M | **13MB** | **394<sub>ms</sub>** | **73MB** | **22<sub>ns</sub>** | 2.7GB | 1.9s | 5.5GB | 117<sub>ns</sub> |
| MHC | 0–250 | 3B | **125MB** | **8<sub>s</sub>** | **41GB** | **53<sub>ns</sub>** | 21GB | 11s | 44GB | 164<sub>ns</sub> |
|  | 150–450 | 3B | **126MB** | **11<sub>s</sub>** | **41GB** | **57<sub>ns</sub>** | 26GB | – | – | – |
|  | 350–650 | 3B | **127MB** | **13<sub>s</sub>** | **41GB** | **55<sub>ns</sub>** | 26GB | – | – | – |
| B.Anth. | 0–250 | 5B | **178MB** | **4<sub>m</sub>22<sub>s</sub>** | **43GB** | **59<sub>ns</sub>** | 34GB | – | – | – |
|  | 150–450 | 9B | **221MB** | **11<sub>m</sub>18<sub>s</sub>** | **43GB** | **59<sub>ns</sub>** | 66GB | – | – | – |
|  | 350–650 | 14B | **254MB** | **23<sub>m</sub>49<sub>s</sub>** | **43GB** | **60<sub>ns</sub>** | 104GB | – | – | – |
| chr22 | 0–250 | 16B | **723MB** | **37<sub>s</sub>** | **34GB** | **60<sub>ns</sub>** | 121GB | – | – | – |
|  | 150–450 | 20B | **737MB** | **52<sub>s</sub>** | **33GB** | **57<sub>ns</sub>** | 149GB | – | – | – |
|  | 350–650 | 21B | **743MB** | **58<sub>s</sub>** | **34GB** | **57<sub>ns</sub>** | 154GB | – | – | – |
| chr1 | 0–250 | 66B | **2.96GB** | **14<sub>m</sub>18<sub>s</sub>** | **43GB** | **63<sub>ns</sub>** | 495GB | – | – | – |
|  | 150–450 | 80B | **3GB** | **45<sub>m</sub>35<sub>s</sub>** | **43GB** | **62<sub>ns</sub>** | 601GB | – | – | – |
|  | 350–650 | 82B | **3GB** | **1h04<sub>m</sub>** | **43GB** | **62<sub>ns</sub>** | 612GB | – | – | – |

## 6 Conclusion

In this work, we have proposed DiVerG, a compressed representation combined with fast algorithms for computing a distance index that exploits the structure of adjacency matrices of sequence graphs to significantly improve space and runtime efficiency. In this way, DiVerG provides a fast solution for the prominent DVP in paired-end read mapping to pangenome graphs. Our extensive experiments showed that DiVerG facilitates the computation of distance indexes, making it possible to solve the DVP when working with large graphs. We have demonstrated how to optimize algorithms with respect to hardware architecture considerations, which has been crucial for processing graphs that capture genomes at the scale of various eukaryotic genomes. We developed DiVerG with a particular focus on aligning paired-end short-reads to graphs, recognizing that the DVP can be a computational bottleneck. In the future, we plan to employ DiVerG in the context of sequence-to-graph alignments.

## Supplementary Material

*Software (Source Code)*: https://github.com/cartoonist/diverg [22] archived at swh:1:rel:362b7551950ee369c01ada82d05b6c8345a3048d

## Acknowledgements

This work was supported by the de.NBI Cloud within the German Network for Bioinformatics Infrastructure (de.NBI) and ELIXIR-DE (Forschungszentrum Jülich and W-de.NBI-001, W-de.NBI-004, W-de.NBI-008, W-de.NBI-010, W-de.NBI-013, W-de.NBI-014, W-de.NBI-016, W-de.NBI-022).

## A Fragment Model in Paired-End Sequencing

Paired-end sequencing read libraries are created by attaching sequencing adaptors at both ends of DNA fragments (inserts), enabling sequencing from both ends. This results in producing two reads per fragment. One read, referred to as *forward read*, extends along the forward strand of the DNA insert. The other one, termed *reverse read* extends from the other end on the reverse complementary strand (Figure 9a).

**Figure 9.**
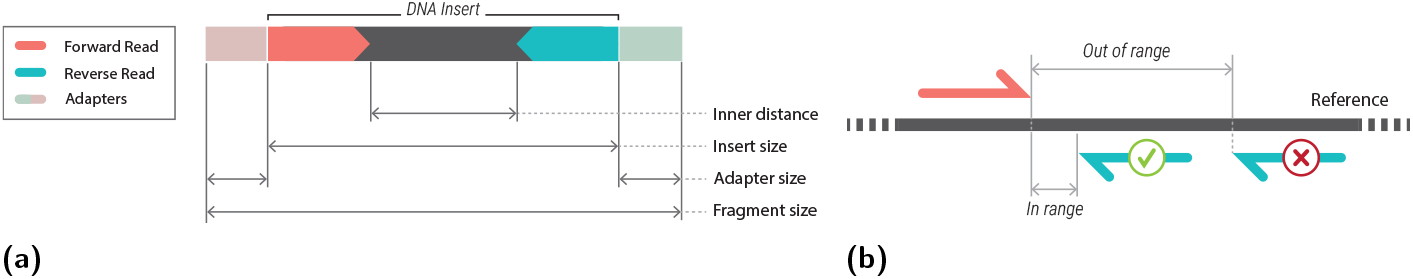
(a) A fragment in paired-end sequencing: a DNA insert with two adapters attached at each end enabling sequencing from both ends. (b) A schematic example of resolving alignment ambiguities using distance validation. The forward read has an unique alignment (in red) which can determine the correct alignment of the other end (blue).

The fragment model of a read library describes the probability distribution of the insert sizes. This model is typically denoted by a normal distribution, characterized by the empirical mean and variance of the insert sizes. In our study, the fragment model is represented by the lower and upper bounds of the expected insert size instead of a probability distribution. These bounds are still probabilistic, as they refer to reasonably chosen cut-offs on the tails of the insert size distribution, e.g. capturing 99.7% of the observed lengths or defined as *µ* ± 3*σ* in which *µ* and *σ* are mean and variance in the fragment model.

There are two ways to define the distance between two paired reads. The *inner distance* between the ends of two paired reads which is determined by the size of unsequenced part of the DNA fragment. It can be calculated by subtracting the sum of the read lengths from the insert size (as shown in Figure 9a). The *outer distance* is defined by the distance between the starts of two paired reads which is equivalent to the *insert size*. Given the variability in both insert sizes and read lengths, and the possibility of overlapping reads in some datasets, using the outer distance can simplify the process of distance validation. Therefore, validating outer distance is more practical compared to the inner distance. Nevertheless, the method is neutral with respect to either definition of distance.

## B Range Compressed Row Storage (rCRS) Analysis

This section explores the rCRS format in more detail, including the construction algorithm and time and space complexity analysis. Before delving into these topics, we present two key observations that follow directly from Definition 6.

### Lemma 13.

*The minimum range sequence of a non-empty set of integers A can be constructed from the sorted array representing A in linear time*.

**Proof**. Construct *ρ*(*A*) by iterating over the elements in *A* in ascending order and including range boundaries: the minimum element, both endpoints whenever there is a gap in the sequence (i.e. *a*_*i*_ ≠ *a*_*i−*1_ + 1 for *i* ≥ 1) and the maximum element. It can be shown that the constructed *ρ*(*A*) has minimum size since any range sequence with smaller cardinality would necessarily imply that it either includes an element not in *A* or misses at least one endpoint in *A*, contradicting Definition 6.

### Lemma 14.

*Given A as a non-empty set of integers, the size of the minimum range sequence of A is at least 2 and at most twice the cardinality of A; specifically*, 2 ≤ |*ρ*(*A*)| ≤ 2|*A*|.

**Proof**. This follows directly from the fact that any range sequence of *A* defines a partition of *A*, denoted as *P*, where each block in *P* represents a contiguous range of integers in *A*. For each block in *P, ρ*(*A*) contains exactly two integers. Thus, 2 ≤ |*ρ*(*A*)| = 2|*P* | ≤ 2|*A*| holds because the number of blocks |*P* | can be at least 1 and at most |*A*|.

### B.1 rCRS Construction

#### Theorem 15.

*rCRS can be constructed from sorted CRS in linear time with respect to the number of non-zero values*.

**Proof**. It directly follows Definition 8 and Lemma 13 (see Algorithm 5).

#### Algorithm 5

Construction of rCRS matrix from sorted CRS.

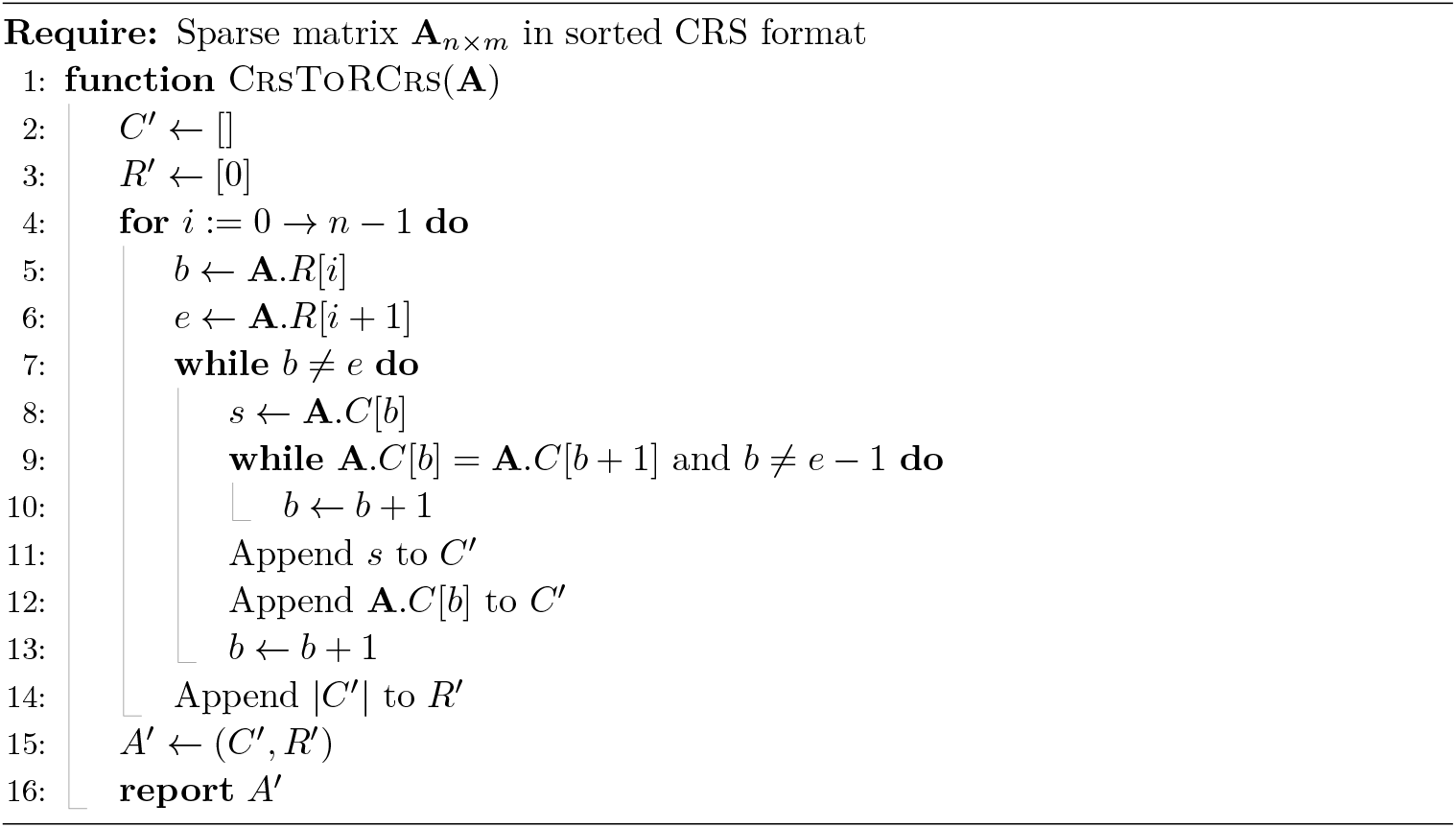

### B.2 Query Time Analysis

#### Theorem 16.

*Let* 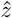 *be the maximum size of the minimum range sequence representation across all rows of sparse matrix* **A**. *Accessing an element in* **A** *represented by rCRS format takes* 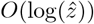 *time*.

**Proof**. By definition, column indices array *C*′ has even size, where each pair of elements at indices 2*k* and 2*k* + 1 forms a range [*C*′[2*k*], *C*′[2*k* + 1]] representing consecutive non-zero values in the corresponding row (Remark 9). Accessing element *a*_*ij*_ requires determining whether column *j* lies within any range in 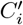, the *i*-th block in the row decomposition of *C*′. Algorithm 6 performs this query using binary search to find the first element in 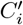 that is greater than or equal to *j* (Line 5). The binary search identifies this element with its index stored in *b*, which is then adjusted to ensure it refers to a range start (Line 13).

Since ranges are sorted and disjoint (Remark 10), only the range [*C*′[*b*], *C*′[*b* + 1]] can potentially contain *j*. The binary search guarantees *C*′[*b* + 1] ≥ *j*, so *a*_*ij*_ = 1 if and only if *C*′[*b*] ≤ *j*. The binary search operates on at most 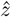 elements, requiring 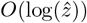 time.

#### Algorithm 6

Querying rCRS matrix.

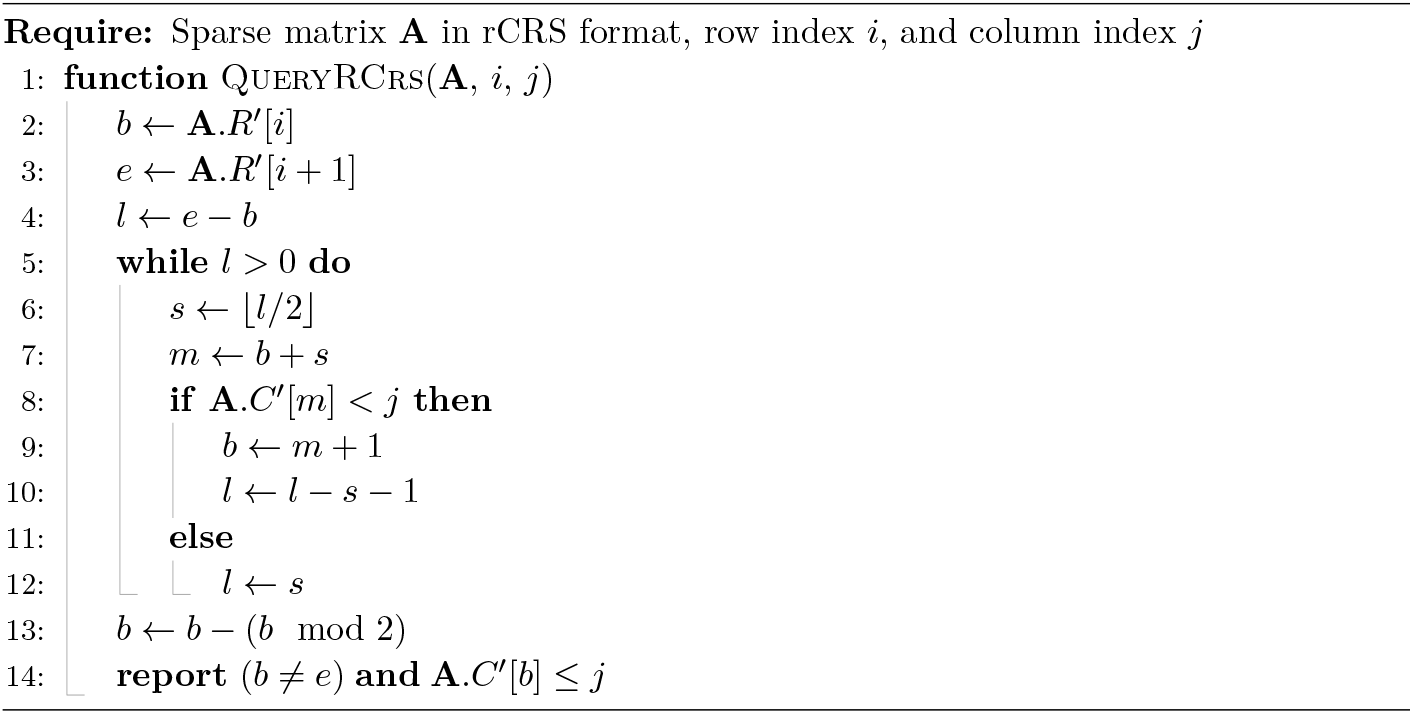

### B.3 Storage Space Analysis

As mentioned before, the memory usage of rCRS format varies according to the distribution of non-zero elements across rows. Since the row map arrays of both rCRS and CRS formats are the same size, only the column indices array *C*′ is the effective factor in compression rate of rCRS compared to the standard CRS.

#### Theorem 17.

*For a sparse matrix A*_*n×m*_, *the column indices array C*′ *in the rCRS representation requires storage space that is at least twice the number of non-empty rows and at most twice the storage needed by the standard CRS format*.

**Proof**. Follows directly from Lemma 14 for each row of *A*.

## C Asymmetrical Range CRS

*Asymmetrical Range CRS* (aCRS) is defined similar to the rCRS format with slight tweaks to avoid redundancy in the column indices array when there is an isolated column index *x*, which is otherwise stored as subsequence [*x, x*] in rCRS.

### Definition 18

(Minimum Asymmetrical Range Sequence). *Let A be a* sorted *sequence of distinct positive integers, and A*_*s*_ *a sequence constructed by replacing any disjoint subsequence of consecutive integers S* = *s*_*f*_ · · · *s*_*l*_ *in A by s*_*f*_ *and −s*_*l*_ *if S >* 1; *otherwise* |*S*| *is replaced by the single element s*_*f*_. *The sequence of minimum size constructed this way is defined as the* minimum asymmetrical range sequence *of A and is denoted by* 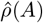.

### Example 19.

Let *A* = [10, 11, 12, 13, 23, 29, 30] from previous example. The minimum asymmetrical range sequence of *A* is defined as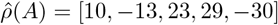.

### Lemma 20.

*The minimum asymmetrical range sequence of a non-empty set of integers A can be constructed from the sorted array representing A in linear time*.

**Proof**. Constructing 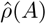 is straightforward. Similar to Lemma 13, it can be done by iterating over the elements in *A* in ascending order and including range boundaries, except we skip adding an endpoint if it equals the last inserted element. Whenever there is an extension, i.e. *a*_*i*_ = *a*_*i−*1_ + 1 for iteration *I* ≥ 1, the next endpoint is negated before insertion. It can be shown that the constructed 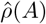 has minimum size since any range sequence with smaller size would necessarily either include an element not in *A* or miss at least one endpoint in *A*, contradicting Definition 18.

### Lemma 21.

*Given A as a non-empty set of positive integers, the size of the minimum asymmetrical range sequence of A is at most equal to the cardinality of A; i*.*e*.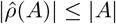.

**Proof**. Similar to Lemma 14, any asymmetrical range sequence of *A* defines a partition of *A*, where each block contributes either one element to 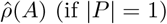 (if |*P*| = 1) or two elements (the endpoints if larger). In either case, each block contributes at most as many elements to 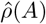 as it contains from *A*. Summing over all blocks shows that 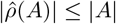.

### Definition 22

(Asymmetrical Range Compressed Row Storage). *For a sparse matrix* **A** *and its sorted CRS representation* (*C, R*), *Asymmetrical Range Compressed Row Storage or aCRS*(**A**) *is defined as two arrays* (*C*^*′′*^, *R*^*′′*^); *where*

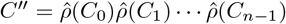, *in which C*_*i*_ *is the i-th block in row decomposition of C; R*^*′′*^ *is an array of length n*+1 *in which R*′(*i*) *specifies the start index in C*^*′′*^ *that corresponds to row i; i*.*e. the start index of* 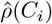. *The last value is defined as R*^*′′*^(*n*) = |*C*′|.

### Remark 23.

Since column indices are non-negative, negative values in aCRS distinctly indicate the upper bound of a range.

### Remark 24.

The elements of the column indices array in aCRS, *C*^*′′*^, are sorted in each block of its row decomposition and the ranges indicated by pairs or isolated elements in 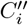 are disjoint.

### Theorem 25.

*aCRS can be constructed from sorted CRS in linear time with respect to the number of non-zero values*.

**Proof**. Directly followed by Lemma 20 and Definition 22.

### Theorem 26.

*For a sparse matrix* **A**, *aCRS*(**A**) *always requires space that is equal to or smaller than that of CRS(***A***) as well as rCRS(***A**).

**Proof**. This can be readily demonstrated using Lemma 21 and Definition 22.

### Theorem 27.

*Let* 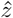 *be the maximum size of the minimum asymmetrical range sequence representation over all rows of sparse matrix* **A**. *Accessing an element in* **A** *represented by aCRS format takes* 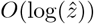 *time in the worst case*.

**Proof**. Each row *i* in aCRS format is represented by a subsequence 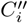 of signed integers (paired or singleton). These elements encode intervals of column indices where non-zero entries occur, and their signs indicate range boundaries (see Remark 23). Similar to rCRS, to query *a*_*ij*_, we perform a binary search on the *absolute values* of 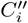 to find the smallest index *b* such that **abs**(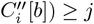. The index *b* is then adjusted to ensure it refers to a range start, based on the sign of 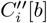 (Line 13). By construction, all ranges in 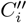 are disjoint and their endpoints are sorted by absolute values (Remark 24), ensuring that at most one range can contain *j* and it is indicated by *b*.

The adjustment step is constant-time, and since binary search performs on 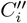 of size at most 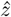, the total time per lookup is 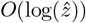. The pseudocode for querying aCRS matrices are given in Algorithm 7.

### Algorithm 7

Querying aCRS matrix.

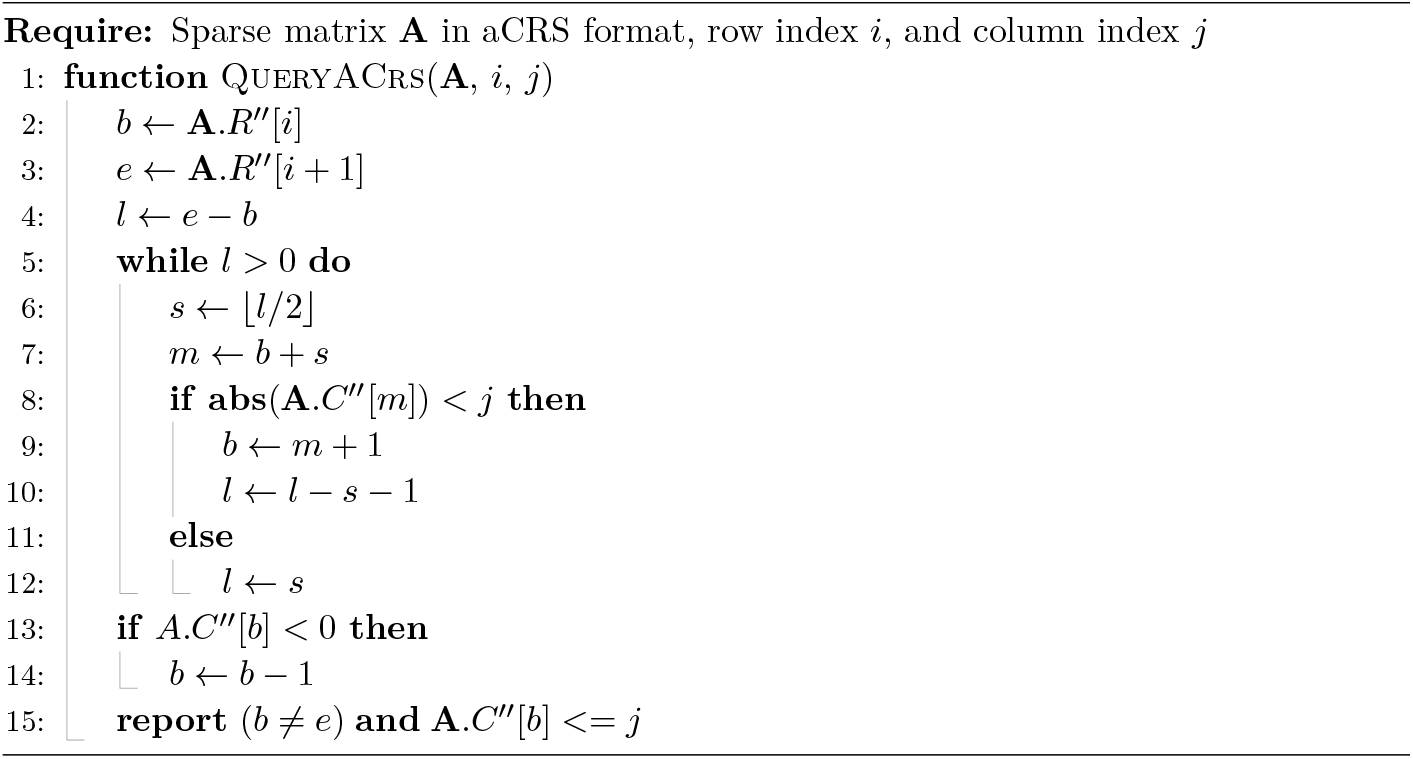

## D Datasets Details

We performed our experiments using various pangenome graphs: mtDNA, BRCA1, LRC_KIR, MHC, B. *anthracis*, and the HPRC CHM-13 pangenome graph for regions chr1 and chr22. This section provides more details about the datasets.

### D.1 mtDNA graph

This graph is constructed using vg [21] and the mitochondrial DNA sequence in GRCh37 reference genome and small variants from the 1000 Genomes project (Phase 3) [1]:

~~~
> vg construct \
        -r human_g 1 k_v 37. fasta \
        -v ALL. chrMT. phase3_callmom - v0_4 .20130502. genotypes. vcf. gz \
        -R MT > \
        ALL. chrMT. phase3_callmom - v0_4 .20130502. genotypes. vg
~~~

### D.2 BRCA1 gene graph

BRCA1 (BReast CAncer type 1) is a gene encoding breast cancer type 1 susceptibility protein. The human BRCA1 gene is located on the long (q) arm of chromosome 17 at region 2 band 1, from base pair 41 196 312 to base pair 41 277 500 (Build GRCh37/hg19). To ensure reproducibility, we re-used the BRCA1 gene graph that was previously published by PairG. It is a variation graph constructed using BRCA1 gene sequence from GRCh37 reference genome and the corresponding variants from 1000 Genomes Project:

~~~
> vg construct \
        -r human_g 1 k_v 37. fasta \
        -v ALL. chr17. phase 3 _shapeit2 _v 5 b .20130502. genotypes. vcf. gz \
        -R 17 :41196312 -41277500 > \
        ALL. chr17 - BRCA1. phase 3 _shapeit2 _v 5 b .20130502. genotypes. vg
~~~

### D.3 LRC_KIR graph

Leukocyte Receptor Complex (LRC) is a locus encoding a large number of immunoglobulin (Ig)-like receptors such as killer cell Ig-like receptors (KIR). In human genome, the LRC region is found at chromosomal region 19q13.4 and spans approximately 1Mbp. Similar to the mtDNA and BRCA1 graphs, this graph can also be constructed using the reference and variants in the 1000 Genomes Project:

~~~
> vg construct \
        -r human_g 1 k_v 37. fasta \
        -v ALL. chr19. phase 3 _shapeit2 _v 5 b .20130502. genotypes. vcf. gz \
        -R 19 :54528888 -55595686 > \
        ALL. chr19 - LRC_KIR. phase 3 _shapeit2 _v 5 b .20130502. genotypes. vg
~~~

### D.4 B. anthracis graph

For this dataset, a de Bruijn graph with a k-mer length of 25 is built using genomes of 20 different strains of B. *anthracis* and splitMEM [32]. The selected 20 strains are based on those reported by PairG and are listed in Table 4. The output of splitMEM is in DOT format and is converted to GFA with a simple Python script we implemented called dot2gfa, which is included with the DiVerG source code:

**Table 4.** List of 20 B. *anthracis* strains used to build the sequence graph and their sizes in Mbp.

| Strain | Assembly id | Size | Strain | Assembly id | Size |
| --- | --- | --- | --- | --- | --- |
| Ames | GCA_000007845.1 | 5.23 | delta Sterne | GCA_000742695.1 | 5.23 |
| SVA11 | GCA_000583105.1 | 5.49 | CNEVA-9066 | GCA_000167235.1 | 5.49 |
| Cvac02 | GCA_000747335.1 | 5.23 | Smith 1013 | GCA_000742315.1 | 5.29 |
| Parent1 | GCA_001683095.1 | 5.23 | 2000031021 | GCA_000742655.1 | 5.33 |
| PR09-1 | GCA_001683255.1 | 5.23 | PAK-1 | GCA_000832425.1 | 5.40 |
| Sterne | GCA_000008165.1 | 5.23 | Pasteur | GCA_000832585.1 | 5.23 |
| 95014 | GCA_000585275.1 | 5.34 | PR06 | GCA_001683195.1 | 5.23 |
| A1055 | GCA_000167255.1 | 5.37 | 55-VNIIVViM | GCA_001835485.1 | 5.35 |
| A0174 | GCA_000182055.1 | 5.29 | Sterne 34F2 | GCA_002005265.1 | 5.39 |
| UT308 | GCA_003345105.1 | 5.37 | Sen2Col2 | GCA_000359425.1 | 5.17 |

~~~
> splitMEM - multiFa - file 20 _strains. fna - mem 25
> dot2gfa. py -k 25 -f 20 _strains. fna -- skip - loops cdg. dot
~~~

### D.5 MHC region graph

A graph for the major histocompatibility complex (MHC) region from reference genome and the corresponding alternative loci reported in GRCh38 [37]. Table 5 lists the sequenced used for construction of MHC graph. This graph, which is constructed based on pairwise alignments using minigraph [30] and seqwish [20], is served as a benchmark for comparing two methods as a mid-sized pengenome.

**Table 5.** The list of alternate loci used for construction of MHC graph.

| NCBI Accession | GenBank Accession | Alternate Locus Group |
| --- | --- | --- |
| NT_167244.2 | GL000250.2 | ALT_REF_LOCI_1 |
| NT_113891.3 | GL000251.2 | ALT_REF_LOCI_2 |
| NT_167245.2 | GL000252.2 | ALT_REF_LOCI_3 |
| NT_167246.2 | GL000253.2 | ALT_REF_LOCI_4 |
| NT_167247.2 | GL000254.2 | ALT_REF_LOCI_5 |
| NT_167248.2 | GL000255.2 | ALT_REF_LOCI_6 |
| NT_167249.2 | GL000256.2 | ALT_REF_LOCI_7 |
| NT_187692.1 | KI270758.1 | ALT_REF_LOCI_8 |

~~~
> minimap 2 -t 20 - cx asm20 *. fa | pigz > mhc - alts. pan. paf. gz
> zcat mhc - alts. pan. paf. gz | fpa drop -l 10000 \
     | pigz > mhc - alts. pan. fpal10k. paf. gz
> seqwish -s mhc - alts. pan. fa. gz -p mhc - alts. pan. paf. gz \
~~~

~~~
     -t 20 -b tmp / -g mhc - alts .pan. fpal10k .gfa
~~~

### D.6 HPRC CHM13 graph

To demonstrate the scalability of the DiVerG in indexing large genome graphs, we employed the CHM13 reference graph released by the HPRC consortium from year 1 data [31]. The graph is built by minigraph [30] with CHM13+Y assembly as the reference ^2^.

## Footnotes

1 http://www-graphics.stanford.edu/~seander/bithacks.html

2 Accessible at: https://github.com/human-pangenomics/hpp_pangenome_resources

